# A nuclear branched-chain amino acid catabolism pathway controls histone propionylation in pancreatic cancer

**DOI:** 10.1101/2025.04.23.650241

**Authors:** Christina Demetriadou, Michael Noji, Austin L. Good, Erick Mitchell-Velasquez, Sharan Venkatesh, Daniel S. Kantner, Jennifer Pennise, Pedro Costa-Pinheiro, Laura V. Pinheiro, Taku Harada, Phuong T.T. Nguyen, Adam Chatoff, Emily Megill, Claudia V. Da Silva Crispim, Mariola M. Marcinkiewicz, Jordan L. Meier, Zoltan Arany, Irfan Asangani, Emma E. Furth, Ben Z. Stanger, Nathaniel W. Snyder, Kathryn E. Wellen

## Abstract

Branched-chain amino acid (BCAA) catabolism contributes prominently to the TCA cycle in the healthy pancreas but is suppressed in pancreatic ductal adenocarcinoma (PDA). The impact of this metabolic remodeling on cancer phenotypes remains poorly understood. Here, we find that the BCAA isoleucine is a primary source of propionyl-CoA in PDA cells. Reduction of propionyl-CoA availability by either genetic perturbation or isoleucine and valine starvation decreases histone propionylation (Kpr) without impacting histone acetylation on specific lysine sites, correlating with reduced transcription of certain lipid- and immune-related genes. Mechanistically, we find that multiple enzymes of isoleucine catabolism unexpectedly localize to and carry out multi-step isoleucine oxidation within the nuclei of PDA cells. Importantly, nuclear localization of the rate-limiting branched-chain alpha ketoacid dehydrogenase (BCKDH) complex is essential for isoleucine-dependent Kpr and gene regulation. Moreover, we demonstrate that isoleucine-sensitive Kpr and its associated gene expression are driven by the MYST family of lysine acyltransferases (KATs), and that the BCKDHA subunit of the BCKDH complex interacts with KAT7 within the nuclear compartment. BCAA catabolism enzymes are apparent in the nuclei of PanIN lesions in mice and PDA tumors in patients, contrasting that in healthy pancreatic acinar and ductal cells. Collectively, these findings unveil a nuclear isoleucine catabolism pathway and highlight its role in controlling histone Kpr and tumorigenic transcriptional programs in PDA.

## INTRODUCTION

Nonmutational epigenetic reprogramming has been described as a hallmark feature of cancer cells (*1*). Histone modifications are a key epigenetic mechanism of transcriptional control and are responsive to the availability of metabolites that serve as substrates of chromatin-modifying enzymes. Well-studied examples include the regulation of histone lysine acetylation (Kac) and histone lysine/arginine methylation by the abundance of acetyl-CoA and S-adenosylmethionine, respectively. However, histones are also decorated by numerous other modifications, whose metabolic regulation and functional significance remain limited (*2*).

Histone lysine propionylation (Kpr) is a three-carbon modification derived from the metabolic intermediate propionyl-CoA (pr-CoA). Though generally less abundant than lysine acetylation, Kpr is a relatively common modification in mouse and human tissues, with certain lysine sites, such as those located on the H3 18-26 peptide, exhibiting propionylation abundance comparable with acetylation (*3*). Kpr has been associated with increased chromatin accessibility, and pr-CoA stimulates transcription in cell-free assays (*4*). Several lysine acetyltransferases (KATs) including p300 (KAT3B), GCN5 (KAT2A), HBO1 (KAT7), and MOZ (KAT6A), have been implicated in mediating Kpr through a high-affinity interaction with pr-CoA (*5–8*). Despite being the second most abundant histone acylation mark in malignant cells, following Kac (*9*), the distinct roles of Kpr and the nutrient sources of pr-CoA that are used by cancer cells for Kpr remain poorly understood.

Recently, we demonstrated that in cancer cells, pr-CoA is highly compartmentalized, with the nucleus showing a distinct enrichment of pr-CoA (*10*). One known metabolic source of pr-CoA that impacts Kpr is the short chain fatty acid propionate, which is produced by the gut microbiota (*11*). In addition to its direct synthesis from propionate, pr-CoA is synthesized from multiple propiogenic pathways, including the catabolism of two of the three branched-chain amino acids (BCAAs), isoleucine (Ile) and valine (Val). These are canonically catabolized in the mitochondria to generate succinyl-CoA that can feed into the TCA cycle (*12*). We found that the nuclear pr-CoA pool in cancer cells was predominately derived from the BCAA isoleucine and feeds into histone propionylation on specific lysine residues, including H3K23 and H4K16 (*10*). These initial findings raised questions about the mechanism of nuclear pr-CoA enrichment and the roles of isoleucine-dependent Kpr in cancer cells.

Upon cellular entry, BCAAs undergo transamination by branched-chain amino acid transaminases (BCATs) to generate branched chain alpha-keto acids (BCKAs), which are then irreversibly decarboxylated by the rate-limiting branched-chain alpha-ketoacid dehydrogenase (BCKDH) complex. In the case of isoleucine catabolism, after multiple other reactions, the acetyl-CoA acyltransferase enzyme ACAT1 catalyzes the formation of pr-CoA and acetyl-CoA from 2-methyl-acetoacetyl-CoA. Pr-CoA is then converted by the propionyl-CoA carboxylase (PCC) into D-methylmalonyl-CoA, and ultimately isomerized to succinyl-CoA, which enters the TCA cycle (*12*) (Fig.1A, schematic).

**Fig. 1.**
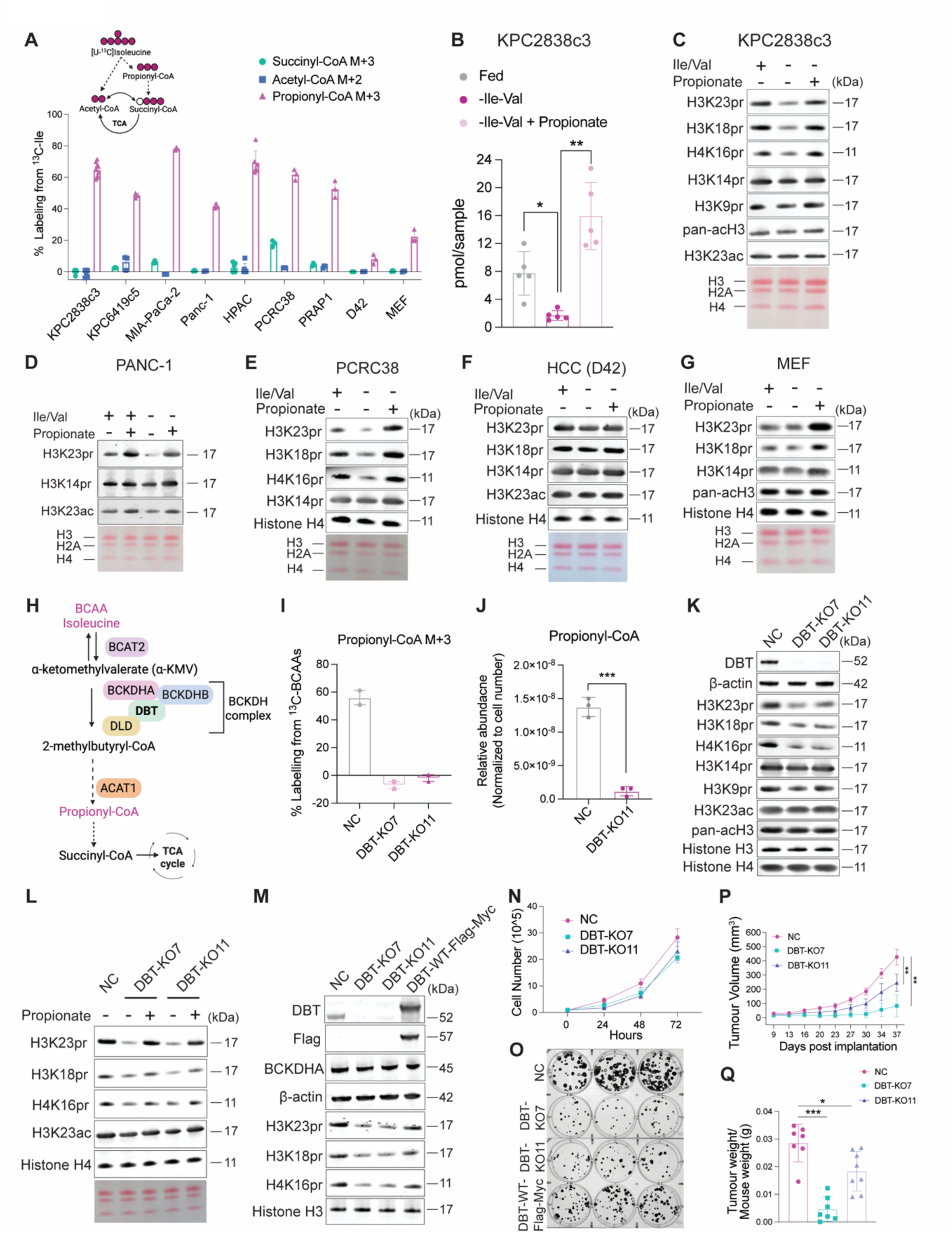
BCAA catabolism regulates histone propionylation and tumor growth in PDA. (**A**) Bar graph showing percent labeling of succinyl-CoA M+3, acetyl-CoA M+2 and propionyl-CoA M+3 from 6 h [U-^13^C]-isoleucine tracing in a panel of cells. The schematic represents carbon labeling contribution from Ile to the indicated CoA metabolites (**B**) Pr-CoA absolute abundance measured by LC-MS after 24 h culture of KPC2838c3 cells in complete media (Fed), starvation (-Ile-Val), or 2 mM propionate rescue (-Ile-Val + Propionate). (**C** to **G**) Western blots of acid extracted histones after 24 h culture +/- Ile and Val, +/- 2 mM propionate in indicated cell lines. Dialyzed serum was used in all experiments to reduce compensation from other metabolites, such as propionate, that are present in the normal calf serum (**H**) Schematic diagram of the BCAA Ile catabolic pathway. (**I**) [U-^13^C]-BCAAs (valine, isoleucine, leucine) were traced into pr-CoA M+3 in DBT-KO (clones 7 and 11) or NC control KPC2838c3 cells. (**J**) Pr-CoA relative abundance measured by LC-MS in whole-cell DBT-KO11 or NC KPC2838c3. (**K** to **L)** Western blots of histones extracted from NC or DBT-KO7/11 cells and treated with propionate. The propionate concentration used for cell treatment was 2 mM unless otherwise specified. (**M**) Western blot in whole-cell extracts or acid extracted histones from NC, DBT-KO7/11 and DBT-KO11 KPC2838c3 cells reconstituted with the human DBT-WT-Flag-Myc DBT protein. (**N**) Proliferation of NC, DBT-KO7, and DBT-KO11 KPC2838c3 cell lines over 3 days of culture. (**O**) Representative images of colony formation assay over 10 days of culture of the indicated KPC2838c3 cell lines. (**P**) Tumor volume of NC and DBT-KO KPC2838c3 cells subcutaneously implanted in C57BL/6J mice. (**Q**) Tumor weight of excised NC and DBT-KO allograft tumors at end point. All experiments were repeated at least twice. Data are presented as mean ± s.d. from independent replicates; Statistical significance was assessed using two-tailed unpaired Student’s *t*-test; *p<0.05, **p<0.01, ***p<0.001.

Pancreatic ductal adenocarcinoma (PDA), an aggressive cancer with a 5-year survival rate of only 13%, has been linked to perturbations in BCAA metabolism. Specifically, elevated plasma BCAA levels, including isoleucine, have been detected in PDA patients years before diagnosis (*13*). BCAA levels are also increased in PDA patient tumors as opposed to adjacent normal pancreas tissue (*14*). Moreover, dietary BCAA restriction or deletion of the pancreatic Bcat2 enzyme in mice confers protection from PDA initiation (*15*). Depletion of key enzymes in the BCAA catabolic pathway, such as BCKDHA, has also been shown to suppress growth in human PDA cells and xenograft tumors, but without impacting mitochondrial metabolism or bioenergetics (*16, 17*). Additionally, BCAA contribution to the TCA cycle is suppressed in PDA tumors (*18*), highlighting the need for further investigation into the mechanistic contributions of BCAA metabolism to tumor progression.

In this study, we investigated the metabolic regulation of histone Kpr and its significance for gene regulation in PDA. We find that pr-CoA is robustly synthesized from isoleucine, despite minimal contribution to succinyl-CoA, across a panel of murine and human PDA cells. Pr-CoA and histone Kpr are highly sensitive to Ile availability, and decreasing pr-CoA abundance by targeting BCAA catabolism enzymes inhibits Kpr, alters gene expression and impairs tumor growth. Remarkably, we find that multiple enzymes of the Ile catabolism pathway localize to the nucleus across PDA cell lines, patient-derived organoids, and patient tumors, and we show that excluding the BCKDH complex from the nucleus suppresses Kpr and Kpr-correlated gene expression. Our findings reveal a previously unknown role of BCAA catabolism in the nuclei of PDA cells and that this compartmentalized pathway is crucial for mediating site-specific histone Kpr and regulating distinct transcriptional programs.

## RESULTS

### BCAA-derived propionyl-CoA is essential for Kpr in PDA cells

We previously found that Ile was a prominent source of pr-CoA in 3 cancer cell lines (11). To more comprehensively assess Ile utilization by PDA cells, we conducted [U-^13^C]-Ile tracing in a panel of murine cell lines (KPC2838c3 and KPC6419c5)(*19*), human established cell lines (MIA-PaCa-2, Panc-1 and HPAC), and human newly patient-derived cell lines (PCRC38 and PRAP1). Ile was indeed a major three carbon supplier for pr-CoA in PDA cells (Fig.1A). In comparison, we observed minor carbon contributions to succinyl-CoA M+3 or acetyl-CoA M+2 isotopologues (Fig.1A), consistent with other major carbon sources contributing to those metabolites. In contrast to the PDA cells, two non-PDA cell lines that we examined, a murine hepatocellular carcinoma (HCC) line D42 (*20*) and mouse embryonic fibroblasts (MEFs) exhibited considerably lower fractional enrichment of pr-CoA M+3 derived from Ile (Fig.1A).

Since Ile catabolism generates pr-CoA, the substrate for Kpr, we next investigated whether Kpr is sensitive to BCAA availability by depriving the cells of isoleucine and valine (Ile/Val) in the culture media for 24 h. Although Val does not contribute to pr-CoA pools in PDA cells (*10*), we removed it along with Ile in our experimental setup to prevent potential compensatory pathways that might arise in the absence of Ile. Ile/Val restriction in KPC2838c3 cells drastically reduced pr-CoA concentration (Fig. 1B), leading to a concomitant decrease in the bulk levels of H3K23pr, H3K18pr and H4K16pr (Fig. 1C). Interestingly, H3K14pr and H3K9pr remained unchanged (Fig. 1C), aligning with prior LC-MS quantification of histone modifications under Ile/Val restriction (*10*). Propionate supplementation, which can generate pr-CoA independently of isoleucine, rescued both pr-CoA and histone Kpr levels (Fig. 1B-C). Proliferation was reduced under Ile/Val restriction and was not restored by propionate, which is unsurprising since Ile and Val are essential amino acids (Supp Fig. S1A). Consistently, Ile/Val deprivation also markedly reduced site-specific Kpr levels in KPC6419c5, PANC-1, and PCRC38 cells, in a manner rescued by propionate (Fig. 1D-E, Supp Fig. S1B). Pr-CoA abundance corresponded with Kpr under these conditions (Supp Fig. S1C). Importantly, unlike Kpr, histone Kac was not responsive to Ile/Val availability (Fig. 1C-D and Supp Fig. S1B), aligning with minimal change in acetyl-CoA abundance (Supp Fig S1C). Contrary to the PDA cells, Kpr was not sensitive to BCAA availability in the murine D42 liver cancer and human 786-O renal cancer cell lines, or in non-malignant MEF cells (Fig. 1F-G and Supp Fig. 1D), suggesting context-dependent regulation.

To confirm that the BCAA catabolic pathway regulates Kpr in PDA cells, we employed CRISPR/Cas9 gene editing in KPC2838c3 cells to delete DBT, the E2 subunit of the BCKDH complex that commits Ile catabolism to pr-CoA production (Fig. 1H and Supp Fig. S1E). Loss of DBT protein in genetically engineered DBT-KO clonal cell lines (DBT-KO7 or DBT-KO11), entirely blocked pr-CoA production from [U-^13^C]-BCAAs compared to non-targeted (NC) control cells (Fig. 1I). Moreover, total intracellular pr-CoA abundance markedly declined in the absence of DBT (Fig.1J). Although acetyl-CoA abundance was also reduced (Supp. Fig. 1F), the percent carbon contribution from [U-^13^C]-BCAAs to acetyl-CoA was markedly lower (∼30-fold) than for pr-CoA (Fig. 1I and Supp Fig. 1G). Consistent with that observed upon modifying nutrient availability, genetic disruption of BCAA catabolism substantially decreased global H3K23pr, H3K18pr and H4K16pr levels (Fig. 1K), which were largely restored by propionate supplementation (Fig. 1L) or the re-expression of a C-terminal Flag-Myc-tagged wild-type DBT protein (DBT-WT-Flag-Myc) in DBT-KO cells (Fig. 1M). In agreement with the Ile/Val deprivation data, the levels of H3K14pr, H3K9pr, and Kac were minimally altered upon loss of DBT (Fig. 1K-L). Additionally, and concordant with DBT deficiency, deletion of BCAT2, the canonically mitochondrial BCAT isoform (Fig. 1H), significantly reduced pr-CoA levels (Supp Fig. S1H) and suppressed Kpr (Supp Fig. S1I). In contrast, deletion of the cytosolic BCAT1 enzyme did not alter histone Kpr (Supp Fig. S1I). Although BCAT2-KO also decreased acetyl-CoA abundance, Kac remained unaffected (Supp Fig. S1H-I). Together, these data demonstrate that the BCAA catabolic pathway is essential for pr-CoA synthesis and site selective Kpr in PDA cells.

We next sought to evaluate the importance of the BCKDH complex in PDA growth. Cell counting over 72 hours showed minimal effect of DBT-KO on cell proliferation (Fig. 1N). Nonetheless, DBT deletion reduced the colony-forming ability of KPC2838c3 cells, which could be partially rescued by exogenous expression of the DBT-WT-Flag-Myc protein, but not with propionate (Fig. 1O and Supp. Fig. 1J). This finding suggests that Ile-derived pr-CoA might promote PDA growth. To assess the impact of DBT-KO on tumor growth *in vivo*, we next subcutaneously injected both KPC2838c3 DBT-KO clones or NC cells into immunocompetent C57BL/6J mice. Consistent with a previous study targeting BCKDHA in PDA cells (15), allograft tumors deficient in DBT exhibited significantly slower tumor growth and reduced tumor weight relative to NC control tumors (Fig. 1P-Q). Although Ki67 expression was not notably different between DBT-KO and NC tumor cells (Supp Fig. S2A), greater infiltration of CD3+ T-cells in DBT-KO tumors was observed (Supp Fig. S2B). Overall, our findings demonstrate that Ile-derived pr-CoA pools influence propionylation at specific histone lysines, and that DBT-dependent BCAA catabolism supports rapid PDA tumor growth.

### The carnitine shuttle pathway is not required for Kpr in PDA cells

The observation of Ile-mediated histone propionylation was surprising, since BCAA catabolism canonically occurs in the mitochondria and acyl-CoA esters cannot cross the mitochondrial membrane directly. The carnitine shuttle pathway is considered the major mechanism for transferring acyl-groups across the mitochondrial inner membrane. In this pathway, mitochondria-generated pr-CoA is converted to propionylcarnitine via the carnitine acetyltransferase (CrAT) enzyme, allowing the propionyl moiety to exit the mitochondrial matrix through the carnitine-acylcarnitine transporter (CAC) (Supp Fig. S3A). This route has been previously implicated in odd chain fatty acid synthesis using pr-CoA as a substrate (*21*), although it is unknown whether it is important for histone Kpr.

Similar to pr-CoA, propionylcarnitine was robustly synthesized from Ile in KPC2838c3 cells but not in D42 cells (Supp Fig. S3B). Furthermore, total cellular propionylcarnitine levels were significantly lower upon Ile/Val restriction in KPC2838c3 and PANC-1 cells (Supp Fig. S3C-D). BCAA starvation also led to a decrease in extracellular propionylcarnitine levels, indicating that propionylcarnitine is present outside of mitochondria (Supp Fig. S3E-F). Additionally, DBT-KO blocked propionylcarnitine synthesis from Ile and decreased propionylcarnitine but not acetylcarnitine abundance (Supp Fig. S3G-H).

Based on these findings, we speculated that shuttling from mitochondria via a propionylcarnitine intermediate is required for BCAA-sensitive Kpr. To test this, we used CRISPR/Cas9 to delete CrAT, CAC, and PCCB, which encodes the beta subunit of propionyl-CoA carboxylase (PCC) (Supp Fig. S3A and S4A). PCC is the mitochondrial BCAA oxidation enzyme that converts pr-CoA to D-methylmalonyl-CoA, and thus its deletion is expected to stimulate propionylcarnitine production. LC-MS analysis revealed reduced propionylcarnitine abundance with CrAT or CAC knockout (Supp Fig. S4B). Consistently, Ile-derived ^13^C incorporation to propionylcarnitine M+3 was strongly inhibited in CrAT-KO and CAC-KO cells (Supp Fig. S4C). Surprisingly, however, Kpr levels were not impaired in KPC2838c3 or PANC-1 cells lacking CrAT or CAC (Supp Fig. S4D-F). Moreover, PCCB deletion did not further boost propionylcarnitine abundance, affect fractional labeling from Ile, or alter Kpr levels (Supp Fig. S4A-D). Under BCAA starvation, Kpr levels were uniformly reduced in both NC and KO cells and were equally restored upon BCAA refeeding (Supp Fig. S4G).

Since initial studies were done in bulk populations, we also generated CrAT and CAC clonal KO cell lines (Supp Fig. S4H-I). LC-MS analysis confirmed a reduction in total abundance and labeling of Ile-derived propionylcarnitine (Supp Fig. S4J-K). However, Kpr levels were unaffected in CrAT-KO and CAC-KO clones under both standard culture conditions and upon Ile/Val refeeding after starvation (Supp Fig. S4L-M). Taken together, these data indicate that although the carnitine shuttle is required for propionylcarnitine synthesis in PDA cells, it is not responsible for supplying pr-CoA to the nucleus for Kpr.

### Nuclear BCAA catabolism regulates Kpr in PDA cells

Since the carnitine shuttle was not required for Ile-dependent Kpr, we considered alternative mechanisms of regulation. Intriguingly, initial immunofluorescence analysis of DBT suggested its presence in the nuclei of PDA cells (Supp Fig. S1E). Previous studies have reported that metabolic reactions typically occurring in the mitochondria can in some contexts also take place in the nucleus. For instance, multi-subunit enzymes of the 2-ketoacid dehydrogenase family, including the pyruvate (PDH) and α-ketoglutarate (α-KGDH) dehydrogenase complexes, have been found in the nucleus in certain contexts, where they supply acetyl-CoA for histone acetylation and succinyl-CoA for histone succinylation, respectively (*22, 23*). Although the mitochondrial BCKDH complex belongs to the same family as PDH and α-KGDH, a distinct nuclear BCAA catabolism pathway has not been described. Moreover, multiple enzymatic steps are required for the synthesis of pr-CoA from isoleucine, and thus a nuclear route to prCoA synthesis would presumably require non-canonical nuclear localization of several enzymes.

To investigate this possibility, we employed a sucrose gradient centrifugation strategy to isolate intact nuclei that were free of whole mitochondria. While the mitochondria-specific protein HSP60 and the cytosolic protein FASN were absent from the nucleus, BCAT2, DBT, BCKDHA, and DLD, were detected in the nuclei of KPC2838c3 cells, along with the nuclear markers lamin A/C and histone H3 (Fig. 2A). Moreover, both the canonical and alternative pr-CoA-producing enzymes, ACAT1 and ACAA1 respectively, were found in the nuclei of KPC2838c3 cells (Fig. 2A). We also validated these findings using immunofluorescence and confocal microscopy. Notably, we observed a distinct nuclear localization of multiple BCAA oxidation enzymes, in addition to their expected non-nuclear presence, while the mitochondrial marker HSP60 was excluded from the nucleus (Fig. 2B, Supp Fig. S1E and S5A). Deletion of DBT reduced the nuclear signal of BCKDHA in both methods, consistent with the formation of a canonical BCKDH complex in the nucleus (Fig. 2A-B). The nuclear presence of DLD-a common subunit of the BCKDH, PDH and α-KGDH complexes-remained relatively unchanged in DBT-deficient cells (Fig. 2-B). Furthermore, the cytosolic BCAT1 protein was not detected in the nucleus (Supp Fig. S5A) in agreement with the observation that BCAT1 loss did not affect Kpr (Supp Fig. S1I). The nuclear localization of BCKDHA, DLD, ACAT1 and ACAA1 was confirmed in the KPC6419c5 cells (Supp Fig. S5B). Conversely, in HCC D42 cells, which do not exhibit BCAA-sensitive Kpr, nuclear localization of BCAA enzymes was not detected (Supp Fig. S5C).

**Fig. 2.**
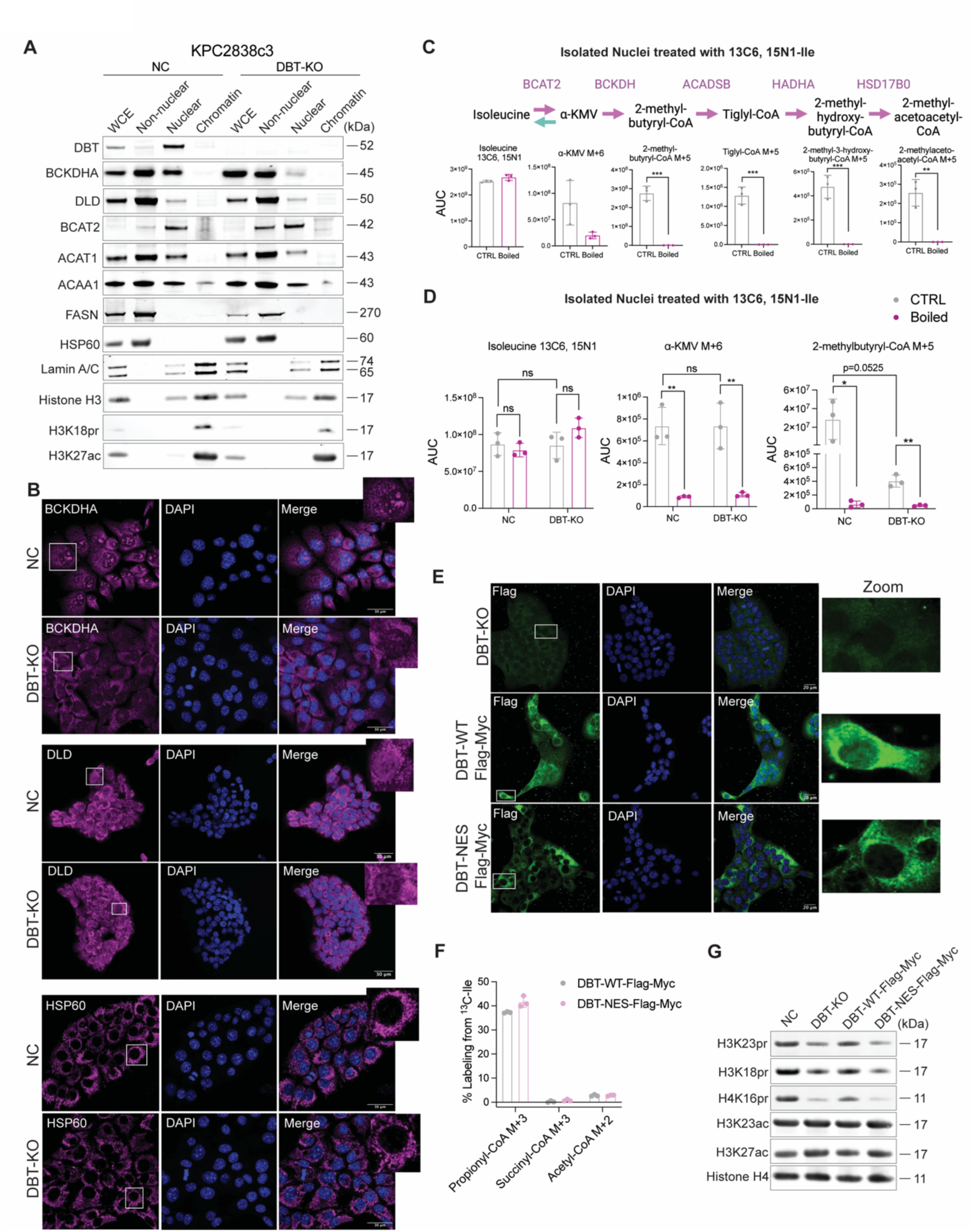
BCAA metabolic enzymes localize to and function in the nuclei of PDA cells. (**A**) Western blot of proteins isolated from the non-nuclear, nuclear, and chromatin compartments of NC and DBT-KO KPC2838c3 cells through biochemical fractionation. Whole cell extract (WCE) was used as a control. DBT-KO11 clone was used in the functional studies unless otherwise indicated. (**B**) Representative immunofluorescence images of the indicated proteins (pink) in NC and DBT-KO KPC2838c3 cells. DNA was stained with DAPI (blue). Scale bars, 30 μm. (**C**) Nuclei were isolated from KPC2838c3 cells and incubated with 13C6, 15N1-isoleucine (Ile) for 18 h, with or without boiling. ^13^C-labeled metabolites of the Ile catabolic pathway were measured and presented as the area under the curve (AUC) in control or boiled nuclear fractions. (**D**) AUC of 13C6, 15N1-Ile, α-KMV M+6, and 2-methylbutyryl-CoA M+5 measured by LC-MS in isolated control or boiled nuclei from NC and DBT-KO KPC2838c3 cells incubated with 13C6, 15N1-Ile for 18 h. (**E**) Immunostaining of Flag tag (green) in DBT-KO, DBT-WT-Flag-Myc addback, and nuclear excluded DBT-NES-Flag-Myc KPC2838c3 cells. Scale bars, 20 μm. (**F**) Tracing of ^13^C carbon labeling to propionyl-CoA M+3, succinyl-CoA M+3 and acetyl-CoA M+2 in DBT-WT-Flag-Myc and DBT-NES-Flag-Myc cells incubated with [U-^13^C]-isoleucine for 6h. (**G**) Histones were extracted from the indicated KPC2838c3 engineered cell lines and analyzed by western blot for the proteins listed on the left of each pane. Data are mean ± s.d. from independent biological replicates. Significance was determined by two-tailed unpaired Student’s *t*-test; *p<0.05, **p<0.01, ***p<0.001.

Although we had previously shown an enriched pr-CoA pool in the nucleus (*10*), we had not examined the intermediate metabolites in BCAA catabolism. To determine whether the detected BCAA catabolic enzymes are catalytically active in the nucleus, we incubated isolated nuclei with ^13^C_6_,^15^N_1_-Ile and monitored the incorporation of labeled carbons into downstream BCAA metabolic intermediates by LC-MS analysis. Intact PDA nuclei efficiently synthesized ^13^C-labeled 2-keto-3-methyl-valerate (α-KMV), 2-methylbutyryl-CoA, tiglyl-CoA, 2-methyl-hydroxybutyryl-CoA and 2-methylacetoacetyl-CoA from Ile (Fig. 2C). Boiling the nuclear fractions abolished this activity, demonstrating that the observed metabolism was enzyme dependent. To conclusively determine whether the BCKDH complex is functional in the nucleus and contributes to 2-methylbutyryl-CoA production, we repeated the isoleucine tracing experiment using intact nuclei isolated from either NC or DBT-KO KPC2838c3 cells. Notably, 2-methylbutyryl-CoA M+5 levels were profoundly decreased in nuclei from DBT-KO cells, while isoleucine and α-KMV levels remained comparable to those observed in NC control nuclei (Fig. 2D). Thus, multi-step isoleucine catabolism enzymes, including the BCKDH complex, are present and active in the nuclei of PDA cells.

To elucidate whether the presence of catalytically active BCAA metabolic enzymes in the nucleus contributes to Kpr, we generated a construct encoding a FLAG-Myc-tagged DBT protein fused to a nuclear export signal (NES) to prevent its nuclear localization (Fig. 2E). Both DBT-WT and DBT-NES constructs expressed similar levels of DBT protein and synthesized comparable amounts of pr-CoA from ^13^C-Ile (Fig. 2F and Supp Fig. S5D). Notably, while exogenous expression of DBT-WT was detected in nuclei and rescued Kpr levels, expression of DBT-NES was excluded from nuclei and failed to rescue Kpr above that in DBT-KO cells (Fig. 2E, G). Collectively, these results demonstrate that multi-step nuclear isoleucine catabolism to pr-CoA supports Kpr through the existence of a catalytically active BCAA catabolic network in the nuclei of PDA cells.

### BCAA oxidation enzymes are present in the nuclei of PanIN lesions and PDA patient tumors

The unexpected presence of the BCAA catabolic pathway in the nuclei of murine PDA cells prompted us to investigate the localization of enzymes of this pathway in mouse genetic PDA models and human PDA samples. Pancreatic tumorigenesis is a multistep process in which pancreatic acinar cells harboring a *Kras* mutation that have undergone acinar-to-ductal metaplasia can give rise to premalignant pancreatic intraepithelial neoplasia (PanIN) lesions (*24*). To explore the localization of BCAA metabolic enzymes during pancreatic cancer development, we performed immunofluorescent analysis on normal pancreatic tissue from WT mice and PanIN lesions from Ptf1α-Cre;Kras^LSL-G12D^ (KC) C57BL/6J mice at 8 weeks (early stage) or 25 weeks (mid stage) of age (Supp Fig S6A). In the healthy WT pancreas, BCKDHA was primarily localized outside of the nucleus of acinar cells (CK19-) and was lowly expressed in normal ductal cells (CK19+). On the other hand, BCKDHA was apparent in the nuclei of PanIN ductal cells at both 8 and 25 weeks of age (Fig. 3A). The DBT protein was similarly detected in the nuclei of *Kras*-mutant PanIN lesions compared to normal ducts (CK19+) or acinar cells (CPA1+) of the WT pancreas, where it was not apparent within nuclei (Supp Fig. S6B). In contrast, the mitochondrial protein HSP60 was strictly detected outside of the nucleus in healthy WT pancreatic cells or PanIN lesions (Supp Fig. S6C). To further validate these findings, we compared PanINs with adjacent normal ducts from 8- or 25-week-old *Kras*-mutant mice. Again, BCKDHA predominately accumulated in the nuclei of PanIN lesions, with minimal detection in the nuclei of adjacent normal ducts (Supp Fig. S7A-B).

**Fig. 3.**
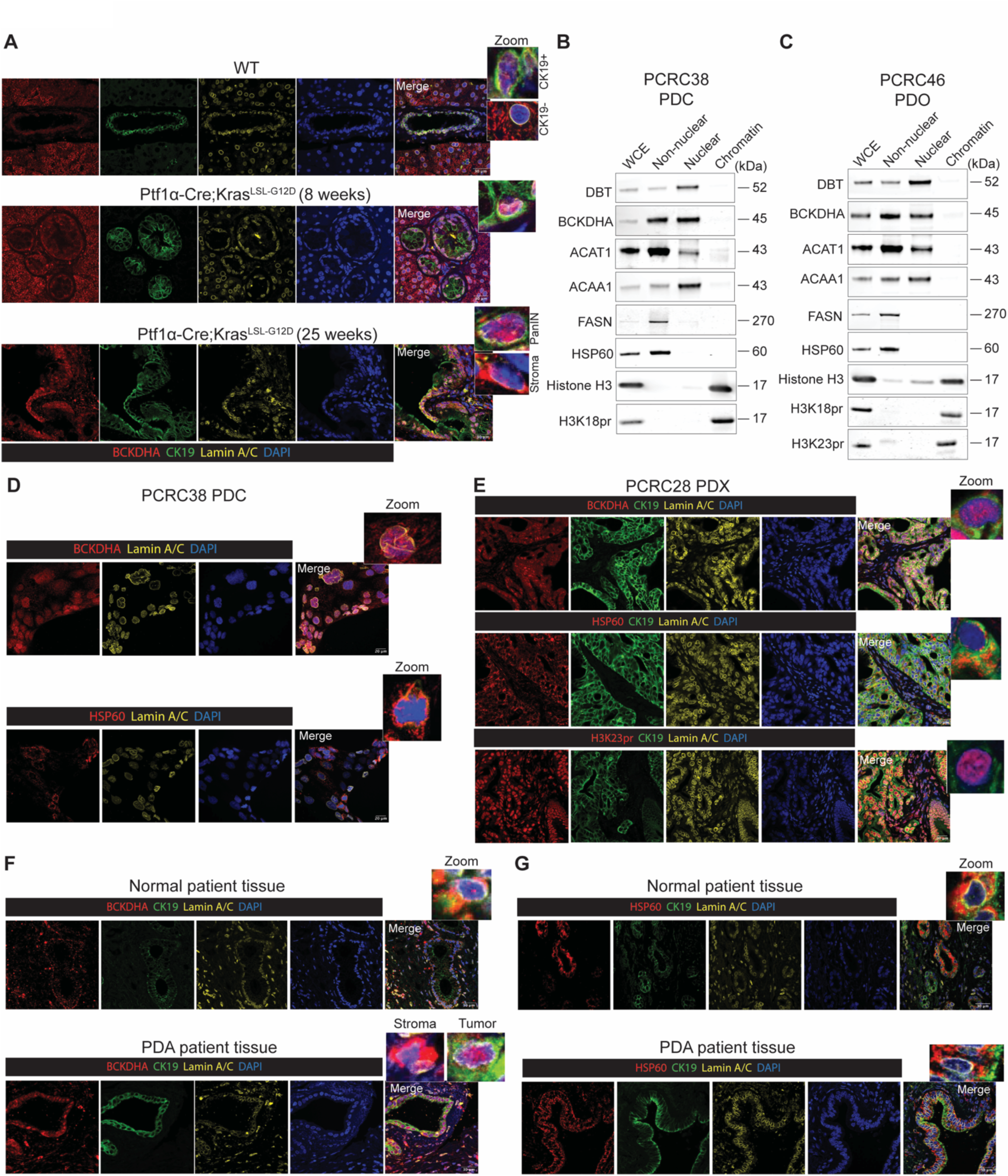
BCAA catabolic enzymes are enriched in the nuclei of PanIN lesions and PDA tumors. (**A**) Immunofluorescence staining of BCKDHA (red), ductal marker CK19 (green) and Lamin A/C (yellow) in the pancreas tissue from WT, 8 weeks old Ptf1α-Cre;Kras^LSL-G12D^, and 25 weeks Ptf1α-Cre;Kras^LSL-G12D^ old C57BL/6J mice. Scale bars, 30 μm. (**B** to **C**) Subcellular fractionation and western blot analysis of (**B**) PCRC38 patient-derived cells and (**C**) PCRC46 patient-derived organoids using antibodies against the indicated proteins. Whole cell extract (WCE) was used as a control. (**D** to **E**) Representative immunofluorescence images of the indicated proteins in (**D**) PCRC38 patient-derived cells and (**E**) PCRC28 patient-derived xenograft tumor. (**F** to **G**) Representative immunostaining images of (**F**) BCKDHA (red) and (**G**) HSP60 (red) in normal and PDAC clinical samples (n=4 biologically independent patients). The CK19 ductal cell marker is shown in green and Lamin A/C control is indicated in yellow color. Nuclei were stained with DAPI (blue). At least 6 areas were imaged in each individual tissue. Scale bars, 30 μm.

We next examined the localization of the BCAA catabolic enzymes in PDA patient-derived samples. Subcellular fractionation analysis identified DBT, BCKDHA, ACAT1, and ACAA1 in the nuclei of patient-derived cell lines grown in 2D culture, as well as 3D patient-derived organoid cultures (Fig. 3B-C and Supp Fig. S8A-B). Furthermore, immunofluorescence staining confirmed the presence of BCKDHA but not HSP60 in the nuclei of patient-derived PDA cells in culture (Fig. 3D). Similarly, unlike HSP60, BCKDHA along with H3K23pr, was robustly detected in the nuclei of patient-derived xenograft tumors (Fig. 3E). Accordingly, in clinical PDA patient samples, BCKDHA was detected in the nuclear compartment of tumor cells (Fig. 3F and Supp Fig. S8C, lower panels), while HSP60 did not exhibit nuclear localization (Fig. 3G and Supp Fig. S8D, lower panels). Conversely, in the healthy pancreas tissue, both BCKDHA and HSP60, were primarily localized outside of the nuclei of normal acinar and ductal cells (Fig. 3F-G, upper panels and Supp Fig. S8C-D, upper panels). Finally, we observed that nuclear localization of BCKDHA was most prominent in PanIN and PDA cells and was not apparent in surrounding stromal cells (Fig. 3A, F and Supp Fig. S7B). Altogether, these data indicate the widespread presence of BCAA metabolic enzymes in the nuclei of PanIN lesions and PDA tumor cells and suggest that nuclear localization of these enzymes is an early event during PDA tumorigenesis.

### Nuclear BCAA catabolism and H3K23pr correlate with regulation of lipid metabolism and immune-related gene expression

Since histone Kpr has been previously associated with transcriptional activation (*4*), we next sought to evaluate the functional significance of BCAA-sensitive Kpr in transcriptional regulation. To this end, we carried out RNA-seq analysis, which revealed distinct clustering between PDA cells cultured in complete media (Fed) and those grown in BCAA-starved (-Ile-Val) media, with or without propionate (Propionate Rescue) (Supp Fig. S9A-B), which restores Kpr but not cell proliferation (Fig. 1K and Supp Fig. S1A). Among the 1,103 downregulated genes in Ile/Val starvation, the expression of 76 genes were rescued by propionate supplementation, whereas none of the 387 upregulated genes were recovered by propionate treatment (Supp Fig. S9C-E). Notably, some of the rescued genes encode lipid metabolism enzymes (e.g. *Acss2*, *Ptges*, *Cyp4b1*, *Mlxipl*, and *Creb3l3*), extracellular matrix, epithelial-mesenchymal transition, and angiogenesis factors (e.g. *Serpinb8*, *Thbs1*, *Grhl2*, *Cthrc1*, and *Fgfbp1*), and immune-related proteins (e.g. *C3ar1*, *Ptges*, *Zbtb32*, *Il13ra1*, and *Ccdc88b*) (Supp Fig. S9F and Supp Table S1). Gene set enrichment analysis (GSEA) of all downregulated genes indicated enrichment in lipid metabolism (fatty acid and cholesterol metabolism) (Supp Fig. S9G).

We also conducted transcriptomic analysis on two DBT-KO clones (DBT-KO7 and DBT-KO11) alongside NC control cells. Principal component analysis revealed a distinct separation between the transcriptional profiles of DBT-KO (DBT-KO7 and DBT-KO11) and NC cells after 48 hours of culture, a condition with minimal impact on cell growth (Supp Fig. S10A-C and 1M). Overall, 302 genes were commonly downregulated, and 105 genes were upregulated in both clonal cell lines (Supp Fig. S10D and S10E). Similar to the Ile/Val deprivation experiments, the downregulated genes mainly included lipid metabolism enzymes (*Ptges*, *Apoa1, Ces1a*, *Vldlr*, *Medag*), drug resistance-associated proteins (*Gstt1*, *Lgi2*, *Sh3rf2*) and immune-related proteins (*Ripor2*, *Ptges*, *Irf6*, *Ceacam1*, *Ly6a*) (Supp Fig. S11A). Conversely, the upregulated genes were enriched for tumor-suppressors and pro-apoptotic proteins such as *Syk*, *Il1rl*, *Bcl2l15*, *Kank1,* and *Crtam* (Supp Fig. S11B). Comparison of the two RNA-seq datasets revealed that 80 genes were commonly downregulated between DBT-deficient and Ile/Val-starved cells (Fig. 4A and Supp Table S1), with 12 of these genes also rescued by propionate treatment (Fig. 4B and Supp Table S1). Pathway analysis of genes downregulated by either Ile/Val starvation or DBT deletion (Kpr-correlated genes) indicated signatures associated with cholesterol biosynthesis, lipid metabolism, drug metabolism, inflammatory signaling, and T-cell response (Fig. 4C). To further examine whether genes are pr-CoA responsive in the setting of DBT deficiency, we performed RT-qPCR analysis in DBT-KO cells treated with or without propionate. As expected, the expression levels of select DBT-responsive genes including *Ptges*, *S100a14*, *Gstt1*, *Ripor2* and *Lgi2* genes were significantly suppressed upon DBT loss in both clones (Fig. 4D). Propionate supplementation in DBT-null cells reversed the low levels of *Ptges*, *S100a14* and *Gstt1*, but failed to restore *Ripor2* and *Lgi2* expression (Fig. 4D), suggesting that specific regulatory pathways may differentially sense Ile-and propionate-derived pr-CoA pools, potentially through spatially controlled mechanisms, in the nucleus.

**Fig. 4.**
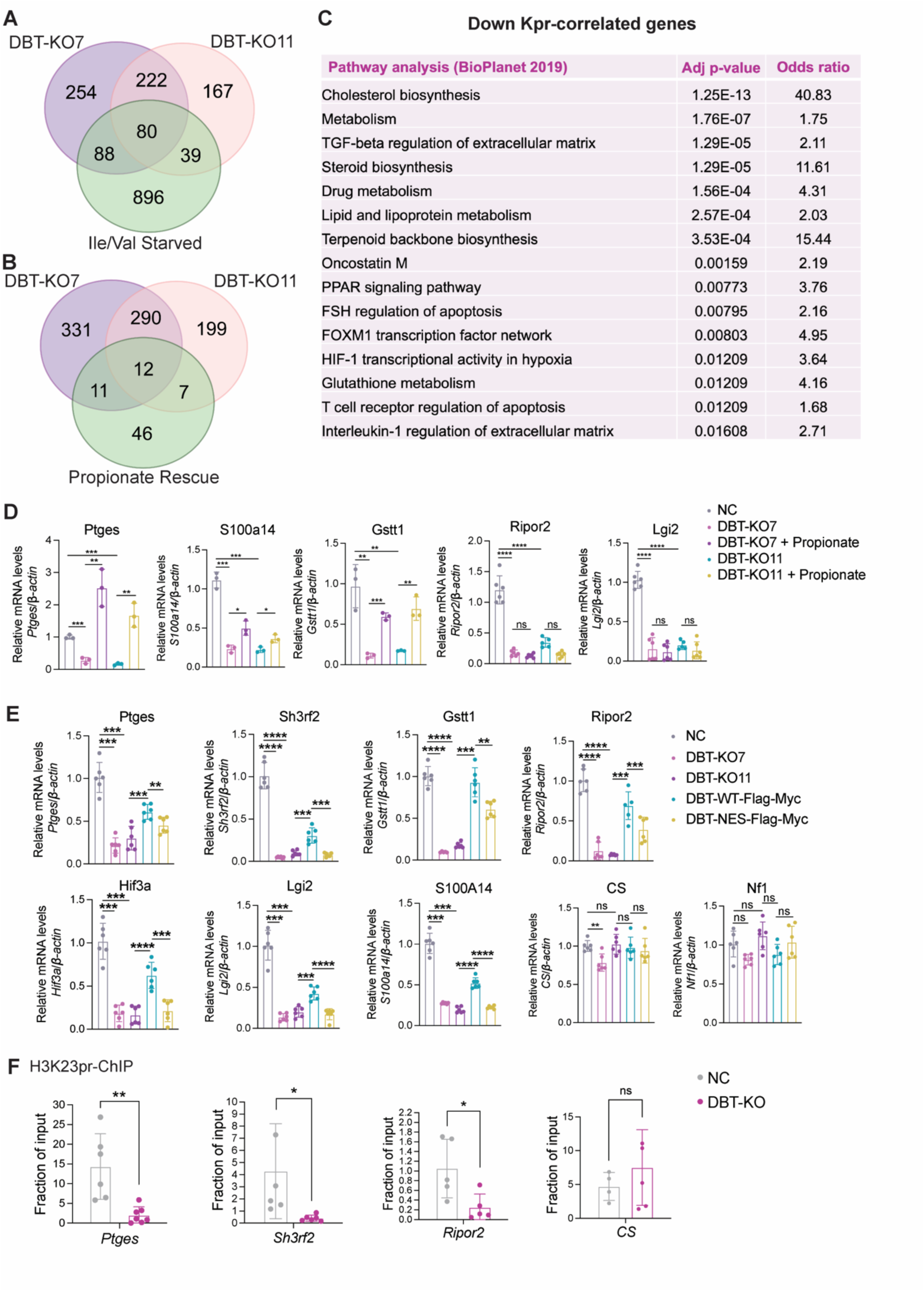
Nuclear BCAA catabolism promotes lipid metabolism and immune related gene signatures in PDA cells. (**A** to **B**) Venn diagrams showing the overlap between downregulated genes in DBT-KO7 and DBT-KO11 with (**A**) BCAA starvation or (**B**) Propionate rescue. (**C**) Pathway analysis of all differentially expressed genes that are downregulated in BCAA starvation and DBT-KO (Kpr-correlated genes). (**D**) qRT-PCR analysis of representative Kpr-correlated genes in NC, DBT-KO7 +/- propionate, and DBT-KO11 +/- propionate cells. (**E**) qRT-PCR analysis of representative Kpr-correlated genes in NC, DBT-KO7, DBT-KO11, DBT- KO11 + DBT-WT-Flag-Myc, and DBT-KO11 + DBT-NES-Flag-Myc KPC2838c3 cells. (**F**) ChIP-qPCR analysis on *Ptges*, *Sh3rf2*, *Ripor2*, and *CS* genomic loci using the H3K23pr antibody. Primers for each locus were designed near the H3K4me3 ChIP-seq peak using the UCSC genome browser. Relative enrichment was normalized to input and the Ct values of rabbit IgG. Data are presented as mean ± s.d. from three independent experiments. Statistical analysis was performed using two-tailed unpaired Student’s *t*-test; *p<0.05, **p<0.01, ***p<0.001, ****p<0.0001, ns=no significance.

Therefore, to interrogate the role of nuclear BCAA catabolism, we leveraged the DBT KO cells expressing either DBT-WT or DBT-NES (Fig. 2F). We found that the reduced transcription of *Ptges*, *Sh3rf2*, *Gstt1*, *Ripor2*, *Hif3a*, *Lgi2* and *S100A14* induced by DBT loss, was significantly rescued by reintroducing DBT-WT protein (Fig. 4E). However, this rescue effect was mostly impaired when exogenously expressing the DBT-NES protein instead (Fig. 4E). In contrast, the expression of control genes *CS* and *Nf1* remained relatively unchanged across different conditions (Fig. 4E). To directly determine if H3K23pr is altered on the promoter of these nuclear BCKDH-regulated genes, we performed chromatin immunoprecipitation (ChIP) followed by qPCR. We found that the promoter loci of *Ptges*, *Sh3rf2* and *Ripor2* were enriched for H3K23pr and that DBT-KO significantly reduced H3K23pr occupancy at these gene loci but not at the *CS* control region (Fig. 4F). Altogether, the results indicate that BCAA catabolism promotes transcriptional regulation of genes involved in lipid metabolism and immune response pathways.

### MYST family KATs regulate nuclear BCKDH-driven Kpr and transcription

The data suggest that Ile-derived pr-CoA is selectively directed to specific lysine sites and genomic regions to regulate transcription. Mechanistically, this specificity could be driven by the consumption of Ile-derived pr-CoA by distinct KAT enzymes that respond to BCAA availability. To identify the KATs responsible for mediating BCAA-sensitive Kpr, we starved PDA cells of Ile/Val and subsequently refed them in the presence of inhibitors targeting different KAT families (*25*). Upon Ile/Val stimulation, H3K23pr was markedly reduced by inhibiting MYST acetyltransferases, using escalating doses of PF-9363 (*26*)(Fig. 5A). Reductions in H4K16pr and H3K14pr were also observed (Fig. 5A). PF-9363 is a potent inhibitor of MOZ/MORF (KAT6A/B)-catalyzed H3K23ac and can also inhibit HBO1 (KAT7)-catalyzed H3K14ac and MOF (KAT8)-catalyzed H4K16ac at higher concentrations (*26, 27*). Degradation or catalytic inhibition of GCN5/PCAF (KAT2A/B) and p300/CBP (KAT3B/A) by GSK702 and CPI-1612, respectively, did not affect these Kpr marks (*28, 29*)(Fig. 5A). Of note, compared to canonical MYST targets (H3K23ac and H3K14ac), H3K23pr appeared slightly more sensitive to MYST inhibition at lower concentrations. Immunofluorescence staining confirmed that MYST inhibition by PF-9363 prominently reduced H3K23pr (Supp Fig. S12A) without changing H3K27ac (Supp Fig. S12B). Conversely, inhibition of p300/CBP by CPI-1612 specifically lowered H3K27ac without impacting H3K23pr (Supp Fig. S12B).

**Fig. 5.**
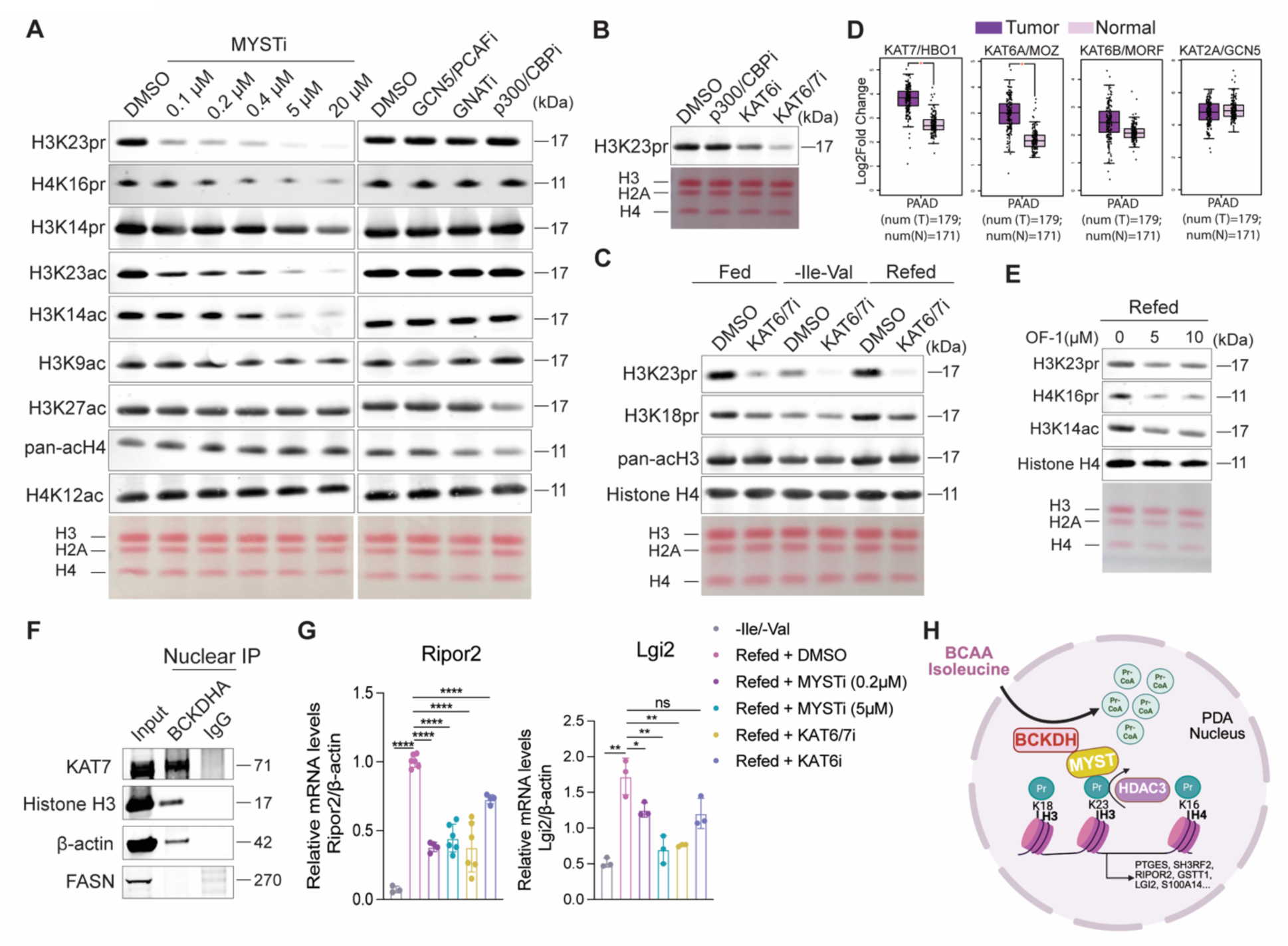
MYST family KATs regulate BCAA-dependent Kpr and associated transcription. (**A**) Western blots of histone proteins extracted from KPC2838c3 cells that were Ile/Val starved for 24 h and then refed for 6 h during which inhibitors targeting the indicated KAT families or DMSO control were added. (**B**) Western blot of H3K23pr in KPC2838c3 cells that were Ile/Val starved for 24 h and then refed for 16 h in the presence of p300/CBPi [CPI-1612] (0.5 μM), KAT6i [WM-1119] (4 μM), and KAT6/7i [WM-3835] (10 μM) inhibitors. (**C**) Western blot of extracted histones from Fed, -Ile-Val (24h), and Refed (16h) KPC2838c3 cells that were treated with KAT6/7i inhibitor WM-3835 (10 μM) or DMSO control. (**D**) Meta-analysis of KAT7, KAT6A, KAT6B, and KAT2A expression levels in PDA patient tissues and matched normal samples from publicly available RNA-seq data extracted from the Gepia data portal (http://gepia2.cancer-pku.cn/#index). (**E**) Western blot of histone proteins extracted from BCAA-refed KPC2838c3 cells treated with the BRPF1/2/3 inhibitor, OF-1, using antibodies against the indicated histone marks. (**F**) Immunoprecipitation of BCKDHA protein or IgG control in isolated nuclei from the KPC2838c3 cells followed by western blot analysis probed with the indicated antibodies. (**G**) qRT-PCR analysis of *Ripor2* and *Lgi2* genes in -Ile-Val and Refed cells treated with the indicated inhibitors or DMSO control. (**H**) Model showing the regulation of Kpr and transcription by the nuclear isoleucine catabolic pathway in PDA. Generated using Biorender. Data are from at least three experiments. Statistical significance was determined by Student’s *t*-test; error is reported as SD (*p<0.05, **p<0.01, ***p<0.001, ****p<0.0001, ns=no significance).

Within the MYST family, KAT6A and KAT7 have been shown to bind pr-CoA with high affinity and exhibit histone propionyltransferase activity *in vitro* and in cells (*5, 8*). In agreement with these studies, selective inhibition of KAT6 (KAT6A and KAT6B), using the WM-1119 inhibitor (*30*), or treatment with the WM-3835, which potently inhibits both KAT7 and KAT6 (*31*), led to a reduction in H3K23pr in BCAA-stimulated PDA cells (Fig. 5B). Supporting these observations, refeeding starved PDA cells with Ile and Val restored H3K23pr and H3K18pr levels, in a manner suppressed by treatment with WM3835 or WM-1119 (Fig. 5C and Supp Fig. 12C). Similarly, depletion of either KAT7 or KAT6A by RNAi in Ile/Val-refed cells suppressed H3K23pr (Supp Fig. S12D). Given the ability of KAT6/7 to impact H3K23pr, we next analyzed the expression levels of these enzymes in PDA patient tumors using TCGA data. Expression of KAT6A and KAT7 was significantly elevated in PDA tissues compared to normal samples, while the levels of KAT6B and KAT2A were not significantly altered (Fig. 5D). KAT6A and KAT7 act within complexes containing BRPF proteins; generally, BRPF2 and BRPF3 interact with KAT7, while BRPF1 forms a complex with KAT6A, to enhance their activity and specificity towards specific chromatin regions (*8, 32–34*). Consistent with involvement of BRPF-containing KAT complexes, targeting the BRPF1/2/3 family in BCAA refed cells using the OF-1 inhibitor suppressed BCAA-sensitive Kpr, along with H3K14ac, which is known to be suppressed by OF-1 (*35*) (Fig. 5E). This result aligns with a previous report demonstrating that BRPF1 is critical for H3K23pr in mouse embryos and fibroblasts (*36*). Together, these results indicate that the MYST family KATs mediate BCAA-sensitive H3K23pr in PDA.

The finding that MYST acetyltransferases mediate specific sites of BCAA-sensitive histone Kpr prompted us to explore whether Kpr is also dynamically regulated by histone deacetylases (HDACs). To test whether HDACs facilitate Kpr removal from histones, we treated PDA cells with general HDAC inhibitors under Ile/Val starvation conditions. Interestingly, only TSA and butyrate, both inhibitors of the HDAC class I and class II enzymes, enhanced H3K23pr levels, whereas inhibition of the HDAC class III (Sirtuins) via NAM or Sirtinol had no effect on H3K23pr compared to the DMSO control (Supp Fig. S12E). Using specific inhibitors, we determined that HDAC class I was responsible for depropionylation at H3K23, H3K18 and H4K16 residues during BCAA starvation (Supp Fig. S12F). To assess the potential clinical relevance of HDAC class I in PDA, we analyzed publicly available RNA-seq data for associations between expression of individual enzymes and overall patient survival. We observed that low expression of HDAC3, but not HDAC1, HDAC2 or HDAC8, correlated with worse survival in PDA patients (Supp Fig. S12G). Moreover, selective inhibition of HDAC3 increased Kpr levels in BCAA-starved cells compared to the DMSO control (Supp Fig. S12H).

Given the observed regulation of Kpr by both nuclear metabolism and epigenetic modifiers, we next investigated potential interactions between BCKDH and nuclear-localized proteins. We performed immunoprecipitation of endogenous BCKDHA from the nuclear fraction, followed by in-gel digestion and mass spectrometry (Supp Fig S12I). In addition to expected BCKDH complex proteins (BCKDHB and DBT), remarkably, KAT7 was also detected in BCKDH immunoprecipitates (Supp Fig. S12J). This interaction was further validated via co-immunoprecipitation followed by western blotting, supporting the ability of nuclear BCKDHA to strongly interact with both KAT7 and chromatin (Fig. 5F).

Finally, to explore whether MYST family KATs are required for regulating Ile-dependent Kpr correlated genes, we treated Ile/Val-refed cells with the MYSTi (PF-9363), KAT6/7i (WM-3835) and KAT6i (WM-1119) inhibitors. While *Ripor2* and *Lgi2* expression increased in BCAA-refed, DMSO-treated cells compared to starved (-Ile-Val) cells, this induction was significantly hindered by all 3 inhibitors of MYST family KATs (Fig. 5G). Overall, these findings demonstrate that BCAA-stimulated Kpr is epigenetically regulated by the opposing functions of KAT6/7 and HDAC3, and that Kpr-correlated gene expression requires MYST family KATs.

## DISCUSSION

In this study, we reveal a multi-step nuclear isoleucine catabolism pathway that regulates histone Kpr and gene expression in PDA cells (Fig. 5H). We demonstrate that PDA cells readily use the BCAA isoleucine as the primary source of pr-CoA. Through nutrient and genetic interventions, we find that histone Kpr at specific lysine residues is highly responsive to BCAA availability and regulated by BCAA oxidation enzymes. Strikingly, we report that BCAA catabolic enzymes, including the canonically mitochondrial BCKDH complex, are present and functional in the nuclei of PDA cells. Furthermore, BCAA catabolic enzymes are detectable in the nuclei of pancreatic premalignant lesions and established PDA tumors in both mice and humans. The nuclear localization of BCKDH is critical for Kpr and the downstream regulation of BCAA-sensitive genes including lipid metabolism and immune response gene signatures. Moreover, we identify MYST family KATs and HDAC3 as the principal BCAA-sensitive epigenetic modifiers responsible for Kpr in PDA cells, and we further show that KAT7 interacts with BCKDHA in the nucleus (Fig. 5H). This study establishes a key link between BCAA catabolism and Kpr in PDA.

Pancreatic cancer remains a deadly disease with a poor prognosis. While prior studies have demonstrated that BCAA catabolism may support PDA tumor growth, mechanistic insights have been lacking, in part because while healthy pancreatic tissue heavily consumes BCAAs for mitochondrial oxidation (*37*), this is suppressed in PDA tumors (*18*). Our data reveal that BCKDH exhibits differential subcellular localization in normal pancreatic tissue compared to PDA tumors, with a pronounced nuclear localization in PDA, raising the possibility that this remodeling of BCAA catabolism participates in tumor progression. These data should prompt further investigation into the significance and mechanisms underlying this reprogramming, as well as whether it could be exploited as a metabolic vulnerability for PDA diagnosis or treatment. Mutations or upregulation of the KAT6/7 acetyltransferases that mediate Ile-dependent Kpr have been linked to oncogenic transcriptional programs (*31, 38*). Recent preclinical and clinical studies have demonstrated promising therapeutic efficacy of newly developed KAT6/7 inhibitors in leukemia and solid tumors (*26, 30, 31, 39*). Given that KAT6A and KAT7 are overexpressed in PDA tumors (Figure 5) and that mutations in KAT6A are observed in PDA (*40*), it would be compelling to investigate the *in vivo* effects of MYST inhibitors on PDA growth.

Recent studies have shown that the epigenome is tightly regulated by the selective import of metabolic enzymes into the nucleus where they support a local supply of metabolic cofactors for chromatin modifications. For example, the nuclear function of acetyl-CoA-producing enzymes such as PDH, ACLY, and ACSS2 in histone acetylation have been increasingly implicated in normal and pathological cellular processes (*22, 41–43*). Comparatively less is known about the existence of nuclear metabolic pathways that regulate other histone acylations and their relevance to diseases including cancer. Nevertheless, a recent study demonstrated that glutaryl-CoA dehydrogenase (GCDH) localizes to the nucleus of glioblastoma stem cells promoting crotonyl-CoA production for histone crotonylation (*44*). Moreover, histone succinylation was found to be regulated by the nuclear presence of the α-KGDH complex in glioblastoma tumor cells (*23*). In this study, we elucidate a nuclear function of the BCAA catabolic pathway in regulating histone propionylation and transcription. These data align with prior proteomics analyses showing that the chromatin-associated protein MED1 interacts strongly with the mitochondrial 2-keto-acid dehydrogenase complexes, including the BCKDH complex, in the nuclei of macrophages (*45*). Additionally, H2A.Z chromatin immunoprecipitation followed by mass spectrometry showed enrichment of isoleucine catabolic enzymes, including DBT, BCKDHA, and ACAT1 proteins in mouse cardiac cells (*46*). These data suggest that nuclear BCAA catabolism warrants further investigation in both physiological and disease contexts.

A key question is what role the nuclear localization of these metabolic enzymes plays, if nuclear pores are permissive to metabolite diffusion. Nuclear localization of metabolic enzymes may facilitate the production of locally derived metabolites from specific nutrients; thus, the spatial distribution of metabolic enzymes within the nucleus may determine nutrient-specific regulation of histone modifications at certain chromatin loci. This specificity appears to depend on histone modifying-enzymes that are found in proximity to these metabolic enzymes. For instance, nuclear GCDH was shown to interact with CBP to drive histone H4 crotonylation from lysine-derived crotonyl-CoA on immunogenic transposable element regions (*44*). Moreover, during CD8^+^ T cell differentiation, glucose- and acetate-derived acetyl-CoA pools are differentially allocated to specific gene loci through the nuclear interaction of ACSS2 and ACLY with p300 and GCN5, respectively (*43*). Interestingly, our gene expression data show that BCAAs and propionate have some overlapping but also distinct effects on Kpr-correlated genes (Fig. 4), suggesting that nuclear pr-CoA pools may be differentially utilized depending on the nutrient source. It could be envisioned that the propionyl-CoA producing enzymes in the BCAA catabolism pathway may interact with distinct KATs than the enzymes that produce propionyl-CoA from propionate. In support of this argument, our study identifies an interaction between BCKDHA and KAT7, implying their potential collaboration in driving Ile-dependent transcription through propionylation on specific lysine sites. This could also explain how H3K14pr, a BCAA-insensitive mark, was shown to be mainly regulated by the GCN5/PCAF acetyltransferases (*4*). As previously reported, BCAT2 and the BCKDH complex form a mitochondrial BCAA metabolon that facilitates metabolite channeling between sequential metabolic reactions (*47*). Given that all metabolites downstream of isoleucine catabolism are being synthesized in the nucleus of PDA cells (Fig. 2C) and that both BCAT2 and BCKDH regulate Kpr and localize in the nucleus (Fig. 1K, 2A and Supp Fig. S1I), future studies should also explore whether Kpr is modulated by the formation of a nuclear-specific isoleucine metabolon localized near chromatin.

The specific role of histone propionylation remains largely unexplored due to the lack of experimental models that can differentially modulate this modification without affecting other histone acylation marks, such as acetylation, since the same writer and eraser enzymes are used for mediating both acetylation and propionylation. Identifying the nuclear BCAA metabolic pathway as a key regulator of Kpr in PDA provides a model that can open new avenues for understanding the distinct functions of this modification in both physiological and disease contexts. Intriguingly, transcriptional analysis from this study indicates that lipid metabolism and immune response genes are key pathways influenced by BCAA-derived Kpr. For example, one of the most responsive targets is *Ptges*, encoding a lipid enzyme responsible for the synthesis of the prostaglandin PGE2, which is secreted by tumor cells to foster an immunosuppressive microenvironment. PTGES has recently been shown to be essential for PDA tumor growth and immunotherapy resistance (*48*). Thus, future studies are needed to elucidate the role of these specific gene targets in promoting PDA progression. Although our data indicate that isoleucine is the prominent source of pr-CoA derived Kpr, BCAA starvation and DBT loss do not eliminate Kpr completely (Fig.1). This suggests that in addition to propionate, other nutrients, such as cholesterol, odd-chain fatty acids, methionine or threonine, may also contribute to pr-CoA pools and Kpr.

Further research is also needed to show how mitochondrial BCAA metabolic enzymes translocate to the nucleus. Different mechanisms have been previously proposed for the translocation of 2-ketoacid-dehydrogenase complexes. The nuclear translocation of the α-KGDH complex is mediated by a nuclear localization sequence in the DLST enzyme (*23*), whereas the PDH complex translocates via MFN2 through tethering of the mitochondria and the nucleus (*49*). However, the specific mechanism governing BCKDH nuclear translocation remains to be identified.

Collectively, our findings uncover a multi-step nuclear BCAA catabolism pathway as the principal metabolic route controlling isoleucine-driven Kpr in PDA. This study enhances our understanding of the role of nuclear metabolism in shaping the cancer epigenome and provides new perspectives on metabolic-epigenetic regulation in cancer biology.

## METHODS

### Cell culture and cell lines

Cell lines were cultured in complete DMEM, high glucose medium (Gibco, 11965084) supplemented with 10% bovine calf serum (Cytiva HyClone, SH30072.04) and 1% penicillin/streptomycin (Gibco, 10378016). PDAC-patient derived cell lines were isolated from biopsy and rapid autopsy, respectively and were cultured in RPMI 1640 medium (10-041-CV). Isoleucine (Ile)- and Valine (Val)-restricted medium was prepared using glucose, glutamine, pyruvic acid and phenol red-free DMEM (Biological Life Science D9800-26) supplemented with glucose (4.5 g/L), glutamine (4mM), non-essential amino acids (2mM), and all essential amino acids except for Ile and Val according to DMEM standard formulation. The pH was adjusted to 7.4 using NaOH and HEPES (25mM). To prevent interference from metabolites such as propionate and propionyl-carnitine present in the serum, 10% dialyzed fetal bovine serum (dFBS) (GeminiBio, 100-108) was used in experiments. The PANC-1 (CRL1469), HPAC (CRL2119), MIA-PaCa-2 (CRL-1420), HEK293T (CRL3216), were purchased from ATCC and MEF (SCRC-1008) cells were generated as previously described (*50*). The murine D42 hepatocellular carcinoma cell line was generated as previously described (*20*). The 786-O kidney adenocarcinoma cell line was gifted by Dr. Celeste Simon lab. The KPC2838c3 (*19*), KPC6419c5 (*19*), PCRC38, PRAP1, PCRC46 and PRAP5 PDAC cell lines were generated and kindly provided by Dr. Ben Stanger. For all Ile/Val restriction experiments, cells were plated at 50%-70% confluency in complete medium with dFBS for 24 hours and were switched to the Ile/Val starvation medium for 24 hours, +/- 2 mM propionate, unless otherwise noted. All cells were grown in a humidified atmosphere at 37°C containing 5% CO_2_ and were routinely tested for mycoplasma through PCR analysis using specific primers (fwd: ACACCATGGGAGYTGGTAAT and rev: CTTCWTCGACTTYCAGACCCAAGGCAT).

### Chemicals and inhibitors

The following HAT inhibitors or protac used in Fig. 5A were gifted from the Dr. Jordan Meier lab: PF9363 (MYSTi) [0.1-20 μM], GSK-4027 (GCN5/PCAF degrader) [1 μM], KAT2i (GNATi) [4 μΜ], and CPI-1612 (p300/CBPi) [0.5 μM]. The WM3835 (HBO1i) [5-20 μM] and WM1119 (MOZi) [1-2 μΜ] inhibitors were purchased from MedChemExpress. TSA (T8552), NAM (HY-B0150), sodium butyrate (303410), and sodium propionate (P1880) were purchased from Sigma. The HDACI (Entinostat), HDACII (MC1568), HDAC2 (CAY-10683), and HDAC3 (RGFP966) inhibitors were purchased from MedChemExpress. For the RNAi experiments, 3×10^4^ cells/ml cells were plated in 6-well plates for 24 h, and 20 nM of siC, siKAT7, or siKAT6A (ON-TARGETplus SMARTpool siRNA, Dharmacon) were added with 12 μl of Lipofectamine RNAiMAX transfection reagent (Invitrogen, 13778150) were added for 48 h.

### Plasmid cloning and generation of engineered cell lines

For generation of stable CRISPR/CAS9 knockout cell lines, sgRNA sequences targeting DBT, CrAT, CAC, PCCB, and negative control (NC) were designed using Benchling (Supp. Table S2) and were cloned into the lentiCRISPRv2 lentiviral vector (*51*). The plasmid sequences were confirmed by Sanger sequencing conducted at the UPenn DNA sequencing core facility. To generate the DBT wild type lentiviral expression vector, human full length DBT (NM_001918) with a C-terminal myc-flag tag was subcloned from the pCMV6 vector (Origene, RC201998) to the pPCP vector (*52*). Nuclear-excluded DBT (DBT-NES) vector was generated by inserting a nuclear excluded localization signal (NES) at the N-terminal of the tag sequence. Plasmid sequences were confirmed by whole-plasmid sequencing. Lentivirus was produced in HEK293T cells using FuGENE HD transfection reagent (Promega, E2311) according to manufacturer’s instructions. The viral-containing media was collected 48 h post-transfection and filtered. Cells were then transduced with the virus in the presence of 10 μg/ml polybrene (Millipore, TR-1003-G) and subjected to 2 μg/ml puromycin (Gibco, 997033-B) or neomycin selection for 72h. For generating clonal cell lines, transduced cells were plated at low density into 96-well plates using serial dilutions to obtain colonies derived from single-cell clones. The efficiency of infection was tested by real-time PCR, western blot and immunofluorescence analysis. Work with recombinant DNA was approved by the Institutional Biosafety Committee (IBC) of the University of Pennsylvania.

### Colony formation assay

Two hundred cells were plated in 6-well plates in triplicates and incubated for 10 days with media changing every 3 days. At the end of the experiment, colonies were stained with 0.5% crystal violet (Sigma, C6158) diluted in 25% methanol in water. Following incubation for 20 min with gentle agitation at RT, cells were washed for 5 times with water and air dried overnight. Images were taken using the Bio-Rad ChemiDoc Touch Imaging System.

### Sub-cellular fractionation

Ten million cells were washed twice with ice-cold 1X PBS and harvested in ice-cold 1X PBS by centrifugation at 1300 rpm for 5 min at 4°C. Cells were resuspended in 250 μl of lysis buffer (10 mM HEPES pH 7.5, 10 mM KCl, 0.1 mM EDTA, 0.5% NP-40, 1mM DTT and 1X protease inhibitor cocktail) and incubated on ice for 20 min with gentle mixing every 5 min. Fifty microliters aliquot (5%) was taken to use as the whole cell extract. Lysed cells were briefly vortexed and centrifuged at 12,000g for 10 min at 4°C. The supernatant was collected as the cytoplasmic (non-nuclear) fraction and pellet was washed 4 times by resuspending in 200 μl of lysis buffer and spun down at 200 g for 5 min at 4°C. The nuclear pellet was then resuspended in 250 μl of Fractionation Buffer (2M sucrose, 1mM MgCl2 and 10mM Tris-HCl pH 7.4) and centrifuged at 16,000 g for 30 min at 4°C. Supernatant was discarded and purified nuclei were washed 2 times by resuspending in 200 μl of lysis buffer and spun down at 200 g for 5 min at 4°C. Purified nuclei were then resuspended in 100 μl of Nuclear Extraction Buffer (20 mM HEPES pH 7.5, 400 mM NaCl, 1 mM EDTA, 1mM DTT and 1X protease inhibitor cocktail) and incubated on ice for 5 min. The sample was centrifuged at 16,000 g for 5 min at 4°C and the supernatant was collected as the soluble nucleoplasm fraction. For isolation of the insoluble chromatin fraction, the pellet was resuspended in 4X laemmli buffer and alternatively boiled at 4°C and chilled on ice 3 times to dissolve. Protein concentration was measured by nanodrop and 20 μg of protein was used for western blotting.

### Stable isotope tracing

For tracing experiments, cells were plated in DMEM containing 10% dFBS. The following day, cells were washed with PBS and tracing media was added for 6h incubation at 37°C and 5% CO_2_. BCAA tracing media was prepared using DMEM base supplemented with glucose (4.5 g/L), glutamine (4mM), non-essential amino acids (2mM), and all essential amino acids except for the targeted experimental tracer. For isoleucine only labeling, [U-13C]-isoleucine (105 mg/L) was added to the tracing media. For all BCAAs labeling, [U-13C]-isoleucine (105 mg/L), valine (94 mg/L), and leucine (105 mg/L) were added. For glucose labeling, DMEM w/o glucose media (Gibco, 11966025) was supplemented with [U-13C]-glucose (4.5 g/L). For unlabeled controls, cells were incubated for 6 h with DMEM supplemented with unlabeled isoleucine (105 mg/L), valine (94 mg/L), leucine (105 mg/L), and glucose (4.5 g/L). For the isotope tracing analysis, FluxFix was used to determine the fractional enrichment into downstream metabolites based on the unlabeled samples (*53*).

### Metabolite extraction and LC-MS

For extraction of acyl-CoAs, culture medium was aspirated from cells in 10-cm plates before adding 1 ml of ice-cold 10% trichloroacetic acid to plates. Plates were scraped on ice to collect cells. Internal standard was added to each sample containing [^13^C_3 15_N_1_]-labeled acyl-CoAs generated in pan6-deficient yeast culture (*54*). Samples were sonicated for 10 x 0.5 s pulses and centrifuges at 16,000g for 10 min at 4 °C. The supernatant was collected and purified by solid-phase extraction using Oasis HLB 1cc (30 mg) solid phase extraction (SPE) columns (Waters). Eluate was dried down under nitrogen gas and samples were resuspended in 50 μl of 5% 5-sulfosalicylic acid (w/v) and transferred to a 96-well plate. Samples were analyzed by an Ultimate 3000 autosampler coupled to a Thermo QExactive Plus instrument in positive electrospray ionization mode as previously described (11). For quantitation, a calibration curve was generated using commercially available unlabeled standards. For extraction of acyl-carnitines, cells were washed with 5ml ice-cold 0.9% NaCl to remove extracellular metabolites and scraped on ice in 1ml −80°C 80% HPLC-grade methanol/20% HPLC-grade water. Extracts were collected in 1.5ml tubes and 10ng of d3-propionyl-L-carnitine internal standard (Cayman 26579) in 50μl 80% HPLC-grade methanol/20% HPLC-grade water was added to each sample. Samples were vortexed and incubated at −80°C for 30 min following centrifugation at 17,000g for 10 min at 4°C. Supernatants were transferred into a 96-well plate and dried under nitrogen gas at room temperature overnight. Dried metabolites were resuspended in 50μl 95% HPLC-grade water/ 5% HPLC-grade methanol using TOMTEC QUADRA 4. For each sample, 1μl was injected and analyzed using a Vanquish Duo UHPLC system coupled to a Thermo QExactive Plus Orbitrap Instrument in positive electrospray ionization mode in full scan mode from 150-1000 m/z. The HPLC system used a hydrophilic interaction chromatography (HILIC) analytical column (Ascentis Express 2.1 mm x150 mm, 2.7μm). The column was kept at 30 °C and the flow rate was 0.5ml/min. The mobile phase was solvent A (10 mM ammonium acetate and 0.2% formic acid in water) and solvent B (10 mM ammonium acetate and 0.2% formic acid in 95% HPLC-grade acetonitrile/5% HPLC-grade water). Elution gradients were run starting from 98% B to 86% B from 0-7 min; 86% B to 50% B from 7-7.3 min; 50% B to 10% B from 7.3-8.3; 10% B was held from 8.3-14.5 min, 10% B to 98% B from 14.5 to 14.510; 98% was held from 14.510-15 min and then the column was equilibrated while eluting on the other identical column. For the analysis of polar metabolites from the nuclear isotope tracing experiment (below), samples were analyzed via a Vanquish Duo UHPLC (Thermo Fisher Scientific) coupled to a Thermo QExactive Plus Orbitrap equipped with a HESI II probe. The samples were injected onto a ZIC-pHILIC 150 × 2.1 mm 5 µm particle size column (EMD Millipore) with a ZIC-pHILIC 20 x 2.1 guard column at a temperature of 25°C. Mobile phases were acetonitrile (solvent A) and 20 mM ammonium carbonate with 0.1% (v/v) ammonium hydroxide in water (solvent B). Initial conditions consisted of a mobile phase composition of 20% B and a flow rate of 0.150 mL/min. A linear gradient program of 20% B to 80% B from 0.5-20 min was performed, followed by a return to initial conditions from 20-20.5 min and an isocratic hold at 20% B from 20.5-28 min. The Orbitrap was operated in polarity switching mode with 70-1000 m/z full scans utilizing an insource fragmentation energy of 1.For each analyte and the internal standard, the peak corresponding to the [M+H]^+^ ion at 5ppm was integrated in Tracefinder 4.1 (Thermo Scientific).

### Nuclear isotope tracing

Intact nuclei were isolated as described above and resuspended in 50 µL of Nuclear Resuspension Buffer (50 mM Tris-HCl, pH 7.5, 0.25 M Sucrose, 25 mM KCl, 5 mM MgCl_2_, and 1X Protease Inhibitor Cocktail). A Reaction Master Mix was prepared containing 1.3 mM NAD, 1.3 mM CoA, 1.3 mM FAD, 1.3 mM α-KG, 46.7 mM MgCl_2_, 1.3 mM TPP, 0.3 mM PLP, 2.7 mM DTT, and 100 mM Tris-HCl, pH 7.5. Each reaction contained 150 µL of Reaction Master Mix and 50 µL of isolated nuclei suspension. The reaction was initiated by adding 2 µL of 100 mM of isotopically-labeled isoleucine (13C6, 15N1-Ile). The reaction was incubated at 37°C overnight (18-20 hours). Boiled controls were prepared by boiling isolated nuclei suspensions at 95°C for at least 5 minutes prior to adding to the Reaction Master Mix. For analysis of polar metabolites (isoleucine and α-KMV), an aliquot of each reaction was added to ice-cold Methanol (final methanol content of 80% v/v) in a 96-well plate. An equal amount of D_3_-propionylcarnitine was added to each sample for use as an internal standard. The samples were dried down under nitrogen and resuspended in 50 µL of 95:5 (v/v) Water:Methanol. The samples were centrifuged for 10 minutes at 2,000 x g and 4°C. Samples were analyzed via HILIC-MS as described above. 1 µL of each sample was analyzed via HILIC-MS as described above. For analysis of acyl-CoAs, an aliquot of each reaction was added to ice-cold 20% trichloroacetic acid (final trichloroacetic acid content of 10% v/v) in a 96-well plate. Acyl-CoA extraction and analysis was performed as described above.

### Protein extraction and western blotting

For protein isolation, cell pellets were washed with 1X PBS and resuspended in lysis buffer (50 mΜ Tris-HCL pH 8.0, 2 mM EDTA, 100 mM NaCL, 1% Triton-X-100, 10% glycerol, 0.5 mM PMSF and 1X protease inhibitor cocktail). Lysis was allowed to continue on ice for 30 min with gentle vortexing every 5 min and lysates were centrifuged at 10,000 rpm for 10 min at 4°C. Total protein concentration was quantified by BCA assay (Thermo Scientific, 2309). For histone acid extraction, cells were lysed in hypotonic lysis buffer (10 mM Tris-HCL pH 8, 1 mM KCL, 1,5 mM MgCl_2_, 0,1% Triton X-100 and 1X protease inhibitor cocktail) and incubated for 30 min on a rotator at 4°C. Nuclei were washed in hypotonic lysis buffer and after centrifugation at 6,500 g for 10 min at 4°C, were resuspended in 0,2 M HCL and incubated overnight with constant rotation at 4°C. Histones were isolated by centrifugation at 6,500 g for 10 min at 4°C and the pH was neutralized with 2 M NaOH at 1/10 of the volume of the supernatant. Thirty micrograms of protein extracts or four micrograms of histone extracts were diluted in 4X laemmli sample buffer (Bio-Rad, 161-0747) containing β-mercaptoethanol and denatured at 95 °C for 5 min. Proteins were separated on gradient SDS-PAGE gels (Invitrogen, NP0322BOX) and transferred to nitrocellulose membranes (Bio-Rad, 1620115). The blots were stained with Ponceau S Staining solution (Cell Signaling, 59803S) and following blocking with 5% BSA in PBST for 1 h at RT, membranes were incubated with primary antibodies in 5% BSA in PBST overnight at 4°C with agitation. Membranes were then incubated with fluorescently labelled secondary antibodies diluted in 5% BSA in PBST for 1 h at RT. All membranes were developed using a LI-COR Odyssey CLx system. Antibodies used in this study were as follows: H3K23pr (PTM-205, PTM BIO; 1:1000), H3K18pr (PTM-213; PTM BIO; 1:1000), H4K16pr (PTM-210, PTM BIO; 1:1000), H3K14pr (PTM-215, PTM BIO; 1:1000), H3K9pr (ABE2852, Millipore, 1:1000), pan-acH3 (06-599, Milipore; 1:1000), H3K23ac (14932S, CST; 1:1000), H3 (ab1791, Abcam; 1:4000), H4 (ab7311, Abcam; 1:1000), H3K27ac (ab4729, Abcam; 1:1000), H3K14ac (7627, CST; 1:1000), H3K9ac (61251, Active Motif; 1:1000), pan-acH4 (06-866, Millipore, 1:1000), H4K12ac (ab177793, Abcam; 1:8000), DBT (PA5-29727, Invitrogen; 1:1000), Flag (F1804, Sigma; 1:1000), BCKDHA (90198, CST; 1:1000), β-actin (3700, CST; 1:1000), CrAT (PAC400Mu01, Cloud-Clone Corp.; 1:1000), CAC (ab244436, Abcam; 1:1000), ACAA1 (12319-2-AP, Proteintech; 1:1000), ACAT1 (16215-1-AP, Proteintech; 1:1000), DLD (HPA044849, Sigma; 1:1000), BCAT1 (TA504360, Origene; 1:1000), BCAT2 (16417-1-AP, Proteintech; 1:1000), FASN (3180, CST; 1:1000), HSP60 (12165, CST; 1:1000), Lamin A/C (2032, CST; 1:1000), and HBO1 (ab70183, Abcam; 1:1000).

### Co-Immunoprecipitation and protein mass spec

Nuclei were isolated as described above. Five percent of the whole cell lysate was kept as an input control. Chromatin in the nuclear fraction was digested with benzonase enzyme (E1014) for 15 min at 37 °C with agitation at 300 rpm followed by sonication (20% amplitude, 10 seconds for two cycles). Sonicated nuclei were centrifuged at 16,000 g for 5 min at 4 °C and the soluble supernatant was diluted to a final volume of 400 μl using the nuclear extraction buffer. Nuclear lysates were mixed with 35 μl of dynabeads Protein A (Invitrogen, 10006D) that were pre-incubated with 8 μg of BCKDHA (Cell Signaling, 12165) or normal rabbit IgG (Cell Signaling, 2729) for 1 h at room temperature with rotation at 18 rpm. Following overnight immunoprecipitation with rotation at 4 °C, the beads were washed 4 times with low salt buffer (10 mM Tris-HCL pH 7.4, 1 mM EDTA, 1 mM EGTA, 150 mM NaCl, and 1% Triton X-100) followed by 1 time in TE buffer (10 mM Tris-HCL and 1 mM EDTA). Immunoprecipitated proteins were eluted in 34 μl of 2X laemmli buffer (BIORAD, 1610747) by boiling at 84 °C for 10 min and separated on an SDS-PAGE gel. For protein mass spectrometry, the SDS-PAGE gel was stained overnight with Coomassie Silver Blue Staining (10% phosphoric acid, 10% ammonium sulfate, 0.12% Coomassie G-250, and 20% methanol) after it was fixed in fixation solution (30% ethanol and 10% acetic acid) for 15 min at room temperature with agitation. Immunoprecipitation in the cytoplasmic fraction was used as a reference control. Trypsin in-gel microdigestion of unique bands detected in the nuclear fraction compared to the cytoplasmic fraction (Supplementary Fig. S12I) and analysis through LC-MS/MS on a Q-Exactive Plus/HF mass spectrometer was performed at the Wistar Proteomics Facility. MS data were searched with full tryptic specificity against the Uniprot mouse database (20230821) and a common contaminant database using Proteome Discoverer version 3.1.1.93.

### Tissue histology

For hematoxylin and eosin (H&E) staining, tissues were fixed in formalin overnight, washed with PBS for 30 min, and dehydrated by titrating in ethanol (50% and 75%) and submitted to the Molecular Pathology and Imaging Core at the University of Pennsylvania for paraffin embedding, sectioning and staining.

### Human tissue procurement for organoid development and imaging

Eligible participants were recruited from clinics at the Hospital of the University of Pennsylvania. Written, informed consent was obtained from patients with pancreatic cancer under Institutional Review Board (IRB)-approved protocols 844610 for tissue collection and organoid generation. For organoid development, tissue samples were minced into small portions and digested at 37°C on a shaker for 1 hour using digestion medium that consisted of 1 mg/mL collagenase XI, 10 μg/mL DNase, 10 μM Y27632, and human wash medium (Advanced DMEM/F-12, 100ug/mL Primocin, 10mM HEPES, 1X GlutaMAX, and 0.1% BSA). Digested cells were washed twice with wash medium and plated in 50 μl domes of Growth-factor Reduced Matrigel (Corning). Organoids were grown in human complete organoid medium containing human wash medium, 500 nM A83-01, 50 ng/mL mEGF, 100 ng/mL mNoggin, 100 ng/mL hFGF10, 10 nM hGastrin I, 1.25 mM N-acetylcysteine, 10 mM Nicotinamide, 1x B27 supplement, RSPONDIN-1 conditioned media (10% final), and WNT3A conditioned media (50% final). For passaging, organoids were dissociated with Accutase, washed twice with human wash medium, resuspended in fresh Matrigel, and expanded in human complete culture medium. 2D cell lines were made from human PDA organoid cultures by recovering cells from the organoid culture that had attached to the plastic bottom of the tissue culture well. The overlying Matrigel domes were dissolved gently with cold human wash buffer, wells were washed with human wash buffer, and cells attached to the plate were expanded in RPMI 1640 medium with 10% FBS and 1X Penicillin/Streptomycin. For imaging analyses, human PDAC tissue was fixed in zinc formalin overnight, transferred to 70% ethanol, and processed for paraffin embedding. Additional tissue sections for imaging were provided by the Cooperative Human Tissue Network (CHTN).

### Immunofluorescence staining

The cells (25×10^4^ cells/ml) were fixed with 4% PFA (J19943-K2, Thermo Scientific) for 10 min at room temperature 24 h after seeding on glass cover slips (15 mm, 1.5 thickness) that were pre-treated with poly-L-lysine (P4707, Sigma). PFA was quenched with 0.1 M ultra-pure Glycine (15527-013) prepared in 1X PBS for 10 min. For immunostaining of histone marks, cells were fixed with 100% ice-cold methanol for 10 min at −20 °C. The cells were then permeabilized with 0.3% Triton X-100 for 10 min. Following blocking in 10% normal donkey serum with 0.5% Triton X-100 in PBS, cells were incubated with the primary antibodies diluted in 5% normal donkey serum with 0.1% Triton X-100 in PBS overnight at 4 °C. The following primary antibodies were used for immunofluorescence: BCKDHA (90198, CST; 1:150), DBT (TA504748, origene; 1:200), DLD (HPA044849, Sigma; 1:300), ACAA1 (12319-2-AP, Proteintech; 1:400), ACAT1 (16215-1-AP, Proteintech; 1:200), HSP60 (12165, CST; 1:400), BCAT1 (TA504360, origene; 1:150), H3K23pr (PTM-205, PTM BIO; 1:100), H3K27ac (ab4729, abcam; 1:400), Ki67 (ab16667, abcam; 1:200), CD3 (ab5690, abcam; 1:100), CK19 (DSHB; 1:200), CPA1 (AF2765-SP, Novus Biologicals; 1:200), Flag (F1804, Sigma; 1:700), and Lamin A/C (ab238303, abcam; 1:800). After washing with PBS for 3 times cells were incubated with secondary antibodies diluted 1:500 in 5% normal donkey serum with 0.1% Triton X-100 in PBS for 1h at room temperature in the dark. The following Alexa fluor secondary antibodies from ThermoFisher Scientific were used: anti-rat Alexa 488, anti-mouse Alexa 555, anti-rabbit Alexa 647, anti-mouse Alexa 647, anti-goat Alexa 488, anti-rat Alexa 555. Nuclei were stained with DAPI (D9542, Sigma). The cover slips were mounted on slides using prolong diamond antifade mounting media (Invitrogen, P36961). For immunostaining of subcutaneous allograft or xenograft tumors, paraffin-embedded tissue sections were incubated at 60 °C for 1 h and rehydrated in the following series of solutions: xylene for 6 min (x2), 100% ethanol for 6 min (x2), 95 % ethanol for 3 min (x2), 70% ethanol for 3 min (x2), deionized water for 5 min, and 1X PBS for 5 min. Antigen retrieval was achieved by boiling the tissue sections in 1X citrate buffer (H-3300) at 96 °C for 20 min. Tissues were allowed to cool down at room temperature for 1 h and washed twice with PBS for 5 min each. Tissue sections were outlined with a hydrophobic pen (Vector Laboratories, H-4000) and permeabilized with 0.3% Triton X-100 for 10 min. Following two washes with 1X PBS for 5 min each, tissues were incubated with blocking buffer (10% donkey serum (D9663, Sigma) diluted in 0.5% Triton X-100 in PBS) for 1 h at room temperature. To reduce mouse antibody reactivity on mouse tissues, one drop of mouse on mouse (M.O.M) blocking reagent (Vector Laboratories, MKB-2213-1) was added to the blocking buffer. Primary antibodies diluted in antibody buffer (1% donkey serum diluted in 0.5% Triton X-100 in PBS) were added overnight at 4 °C. The next day tissues were washed three times with PBST (0.1% Tween in 1X PBS) for 5 min each and secondary antibodies diluted in antibody buffer were added for 1 h at room temperature. Nuclei were stained with DAPI. The slides were mounted using prolong diamond antifade mounting media. For immunostaining of patient tissues, paraffin-embedded tissue sections were processed similarly with the following alterations: Tissues were rehydrated in xylene for 6 min (x3), 100% ethanol for 6 min (x2), 95 % ethanol for 4 min (x2), 70% ethanol for 4 min (x2), deionized water for 5 min, and 1X PBS for 5 min. Antigen retrieval was achieved by boiling the tissue sections in 1X citrate buffer at 96 °C for 30 min. Imaging was carried out using a ZEISS LSM 880 or Leica STELLARIS confocal microscope and confocal images were processed using ImageJ.

### RNA extraction and RT-qPCR

Total RNA was extracted from cells using the RNeasy Mini kit (Qiagen, 74104) and was treated with DNAse (Qiagen, 79254) according to the manufacturer’s instructions. An amount of 0.5 μg total RNA was then reverse transcribed to cDNA using the high-capacity RNA-to-cDNA master mix (Applied Biosystems, 4368814). cDNA was diluted 1:20 and amplified with PowerSYBR Green PCR Master Mix (Applied Biosystems, 4367659) using a ViiA-7 Real-Time PCR system. Fold change in expression was calculated using the ΔΔC_t_ method and expression data were normalized to the β-actin and GAPDH housekeeping genes. Primer sequences were obtained from Sigma and are listed in Supplementary Table S2.

### RNA-sequencing

Total RNA was isolated using the RNeasy Plus Mini Kit (Qiagen, 74134) following the manufacturer’s protocol. RNA integrity was assessed using the 4200 TapeStation system (Agilent) in combination with the RNA ScreenTape Analysis Kit (5067-5576), according to the supplier’s instructions. Poly(A)+ RNA was isolated from total RNA using the NEBNext Poly(A) mRNA Magnetic isolation module (E7490S). RNA-seq libraries were generated using the NEBNext® Ultra™ II Directional RNA Library Prep Kit for Illumina (NEB, E7765S). Library quality and fragment size were evaluated on the Agilent TapeStation system (G2991BA) using the Agilent High Sensitivity D1000 ScreenTape (5067-5584) prior to sequencing on the Illumina NextSeq 2000 platform. Reads from fastq files were mapped to the mouse genome (GRCm38) using Salmon (*55*) with ‘libType A’ and ‘validateMapping’ options. The reads mapped to coding genes were then used as an input to DESeq (*56*) to perform differential analysis. Differential gene expression was determined at a threshold of p < 0.05 and fold change > 1.5. Volcano plots from the DESeq results were plotted using the EnhancedVolcano package (DOI: <u>10.18129/B9.bioc.EnhancedVolcano</u>) and PCA plots were plotted using the plotPCA package. Venn diagrams for the commonly upregulated and downregulated genes were generated using the ggVenn package or the https://bioinformatics.psb.ugent.be/webtools/Venn/ tool. For the pathway enrichment analysis, the gene counts were used as input to the GSEA program (*57*). Enrichment analysis was performed on the mouse hallmark gene sets from MSigDb (*57*). Heatmaps were generated using the normalized counts from DESeq using the pheatmap package (*58*). In addition, ashr (*59*), ggplot2 (*60*), snakemake (*61*), tximport (*62*), and rtracklayer (*63*) packages were used for performing these analyses. Pathway analysis was performed using the Enrichr web server (*64*).

### Chromatin immunoprecipitation

Cells in 15 cm plates were crosslinked in 1% formaldehyde for 10 min, quenched with 125 mM of glycine for 10 min at room temperature, scraped in 1X ice-cold PBS and collected by centrifugation at 1600 rpm for 5 min at 4 °C. After the cells were lysed in SDS lysis buffer (1% SDS, 10 mM EDTA, 50 mM Tris-HCL pH 8, and 1X protease inhibitor cocktail), samples were sonicated (30 sec ON/30 sec OFF for 13 cycles) in a Bioruptor (Diagenode) to obtain chromatin fragments between 250-600 bp. The chromatin was collected by centrifugation at 13,000 rpm for 10 min and diluted 1:10 in IP buffer (1% Triton X-100, 2 mM EDTA, 50 mM Tris-HCL pH 8, 150 mM NaCl, and 1X protease inhibitor cocktail). The appropriate amount of antibody was incubated with 40 μl of dynabeads Protein A (Invitrogen, 10006D) for 1 h at room temperature. Twenty-five micrograms of chromatin were incubated with the antibody-beads complex for 5 hours at 4 °C with rotation, while 10% of the diluted chromatin used for immunoprecipitation was kept aside for the 10% input. The immunoprecipitates were then washed as follows: two washes with low salt buffer ( 0.1% SDS, 1% Triton X-100, 2 mM EDTA, 20 mM Tris-HCL pH 8, 150 mM NaCl, and 1X protease inhibitor cocktail), 1 wash with high salt buffer: 0.1% SDS, 1% Triton X-100, 2 mM EDTA, 20 mM Tris-hcl pH 8, 500 mM NaCl, and 1X protease inhibitor cocktail), two washes with low salt buffer and one wash with TE buffer (10 mM Tris-HCL and 1 mM EDTA). Immunoprecipitated chromatin was eluted from the beads by incubating the samples in 200 μl of freshly prepared elution buffer (1% SDS, 0.1 M NaHCO3) at 950 rpm for 30 min at 25 °C. Next, cross-linking was reversed by incubating the eluted chromatin and 1% input (diluted in elution buffer) with 200 mM NaCl containing 0.5 μg/μl RNase (AM2271, Invitrogen) at 65 °C overnight with rotation (300 rpm). DNA was purified using a Qiaquick PCR purification kit (Qiagen, 28106) and eluted in 50 μl DNase/RNase free water. ChIP was analyzed by RT-qPCR as described above and 2 μl of DNA was used in each reaction. Primer sequences were obtained from Sigma and are listed in Supplementary Table S2.

### Tumor allograft models

To generate allograft models, 300,000 KPC2838c3 cells (NC, DBT-KO7, DBT-KO11) were resuspended in 100 μl of serum-free DMEM medium and injected subcutaneously into the right flank of 6-week-old C57BL/6J male mice purchased from the Jackson Laboratory (000664) (n=7/group). Before the injections, mice were allowed to acclimate for 7 days at the animal facility of the Biomedical Research Building at UPenn. All mice were housed in standard conditions, with five animals per cage. Throughout the experiment, mice were supervised for their overall health condition. Tumor volume was monitored every 3-4 days using a digital caliper. At the end of the experiment, mice were euthanized, and tumors were excised, weighted, and processed for histology. All animal work was approved by the Institutional Care and Use Committee (IACUC) of the University of Pennsylvania.

### Patient-derived tumor cell implantation and harvesting

PDAC organoids were passaged as described above, resuspended in Matrigel, and kept on ice until injection. Tumor cells (1 × 10^5^ to 1 × 10^6^) were injected subcutaneously into 6- to 8- week-old NSG mice. After removal, tumors were washed 3× with PBS. Tissues were then fixed overnight in zinc formalin. After 24 hours, tissues were transferred to 70% ethanol and processed for paraffin embedding. Tissues were sectioned at 5 μm.

### Statistical analysis

Graphpad Prism software (v.10) was used for graphing and statistical analysis. Data presented are shown as mean ± s.d. For comparisons between two groups, datasets were analyzed by two-tailed unpaired Student’s *t*-test and statistical significance is defined as p < 0.05 (*), p < 0.01 (**), p < 0.001 (***), p < 0.0001 (****), ns=no significance.

## ACKNOWLEDGMENTS

We would like to thank all Wellen and Snyder lab members for thoughtful discussion; the Wistar Proteomics Facility for assistance and analysis with the proteomic studies; the Penn Molecular Pathology and Imaging Core for tissue histology; Nivitha Murali, Romie Azor, and Alison Jaccard for their assistance. This research was supported by a Ludwig Institute for Cancer Research grant to K.E.W., NCI grants R01-CA-174761 to K.E.W., R01-CA-248315 to K.E.W and Z.A, and R01-292937 to K.E.W. and N.W.S. C.D. was supported by the AACR Anna D. Barker Basic Cancer Research Fellowship. Funding support for The Wistar Institute core facilities was provided by Cancer Center Support Grant P30CA010815. J.L.M. was supported by the Intramural Research Program of the National Institutes of Health (NIH), the National Cancer Institute, and The Center for Cancer Research (ZIA BC011488). M.N. was supported by NCI grant F31CA261041. P.C.P. was supported by K00CA245802. L.V.P. was supported by T32DK007314 and a diversity supplement to R01CA262055. B.Z.S. was funded by NCI grant R01-CA-252225, the Penn Pancreatic Cancer Research Center, and A Love for Life.

## AUTHOR CONTRIBUTIONS

Conceptualization: C.D., K.E.W., and N.W.S. Methodology: C.D., A.L.G., E.M.V., D.K., J.L.M., and N.W.S. Investigation: C.D., M.N., A.L.G., E.M.V., D.S.K., J.P., P.C.P., L.V.P., T.H., P.T.T.N., A.C., E.M., and C.V.D.S.C. Formal analysis: C.D., S.V., D.S.K, A.C., E.M., C.V.D.S.C., E.E.F, and N.W.S. Writing-original draft: C.D., K.E.W., and N.W.S. Writing-review and editing: C.D., M.N., A.L.G., E.M.V., S.V., D.S.K., J.P., P.C.P., L.V.P., T.H., P.T.T.N., A.C., E.M., C.V.D.S.C., J.L.M., Z.A., I.A., E.E.F., B.Z.S., N.W.S., and K.E.W. Supervision: K.E.W., and N.W.S. Funding acquisition: K.E.W., and N.W.S.

## DATA AVAILABILITY

RNA-seq data generated in the paper are deposited in Gene Expression Omnibus under accession number GSE294027.

