## Supplementary figures and images for "A nuclear branched-chain amino acid catabolism pathway controls histone propionylation in pancreatic cancer"

### S1

##### Supplementary Figure S1

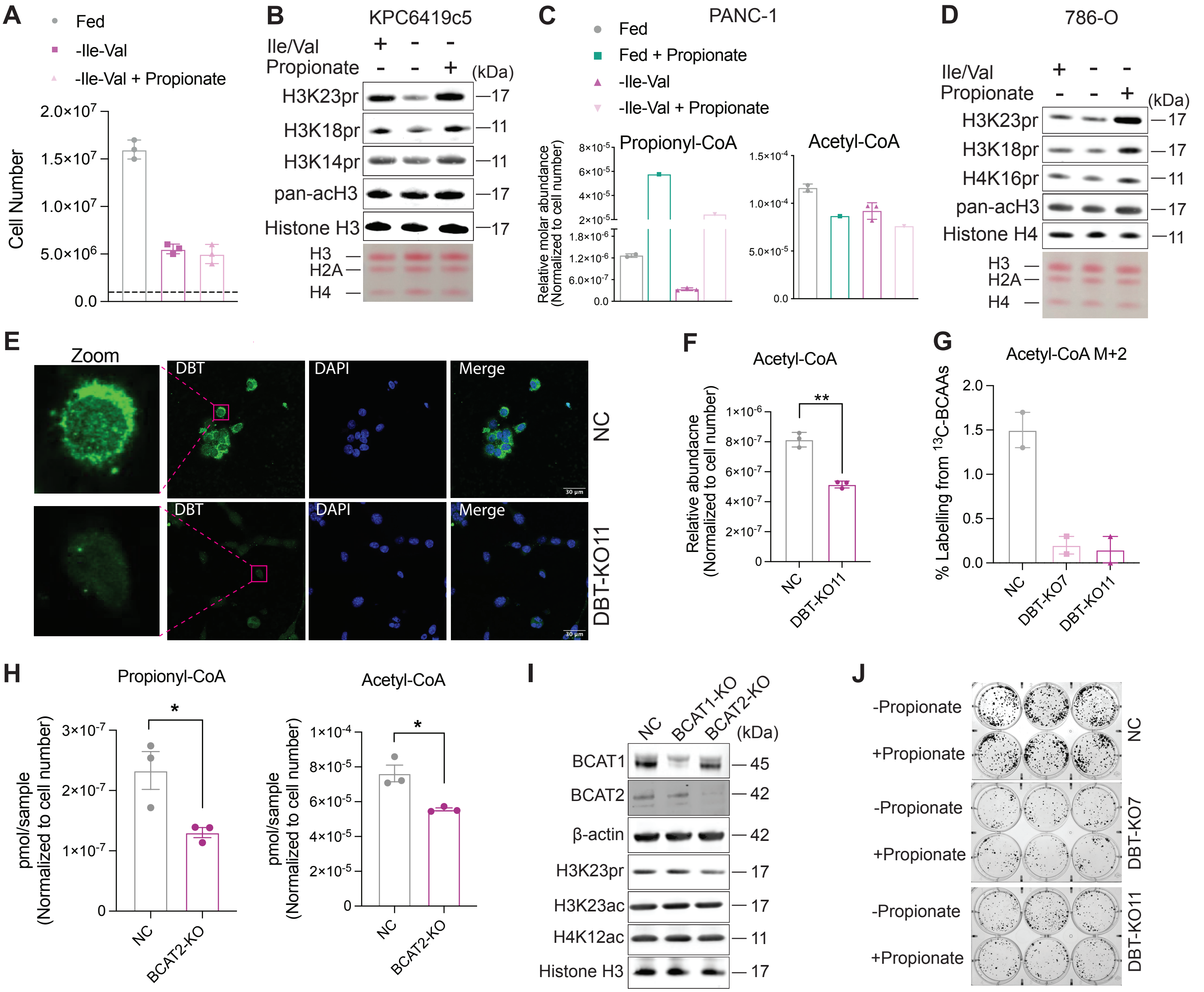

### S2

##### Supplementary Figure S2

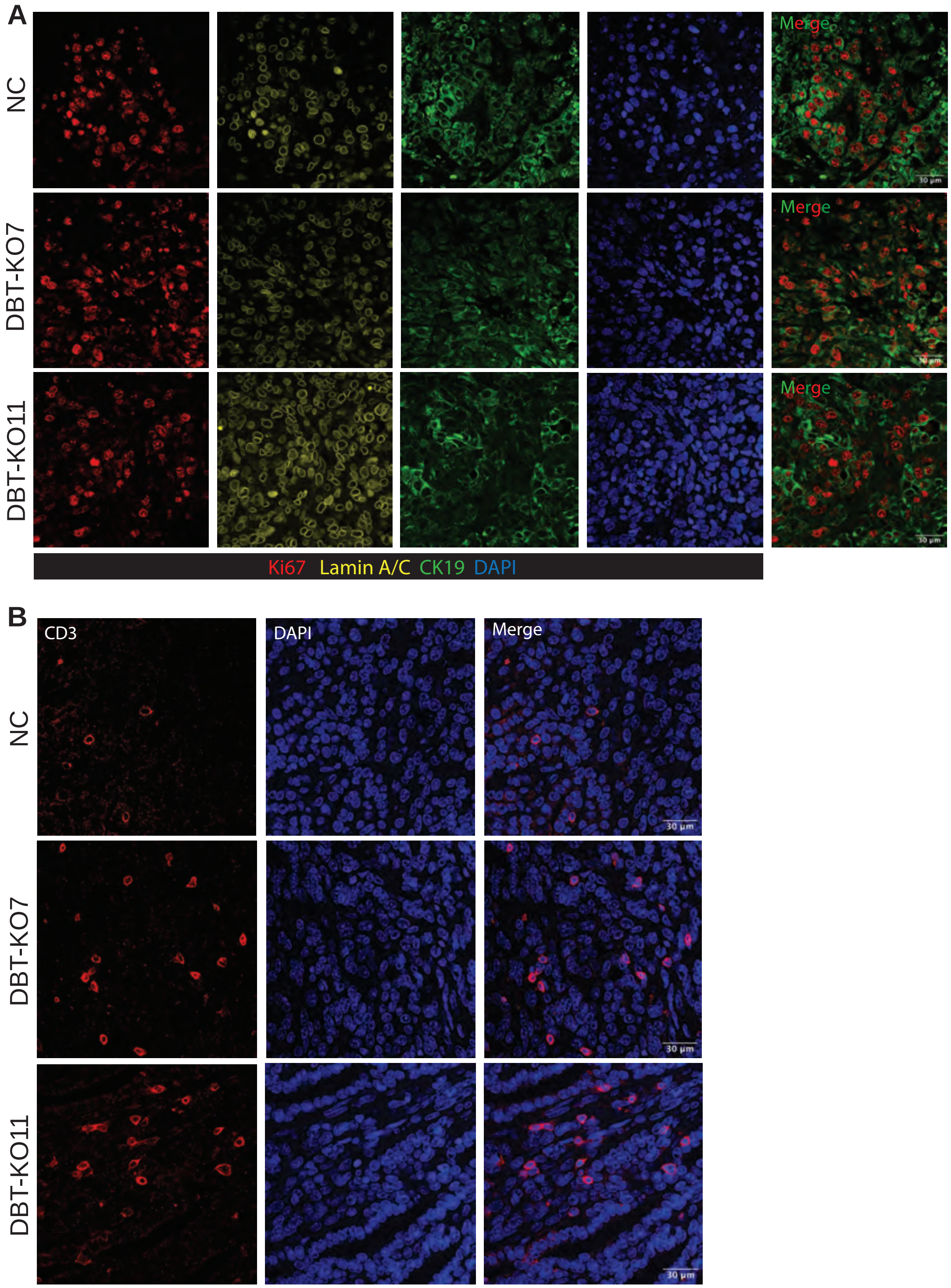

### S3

#### Supplementary Figure S3

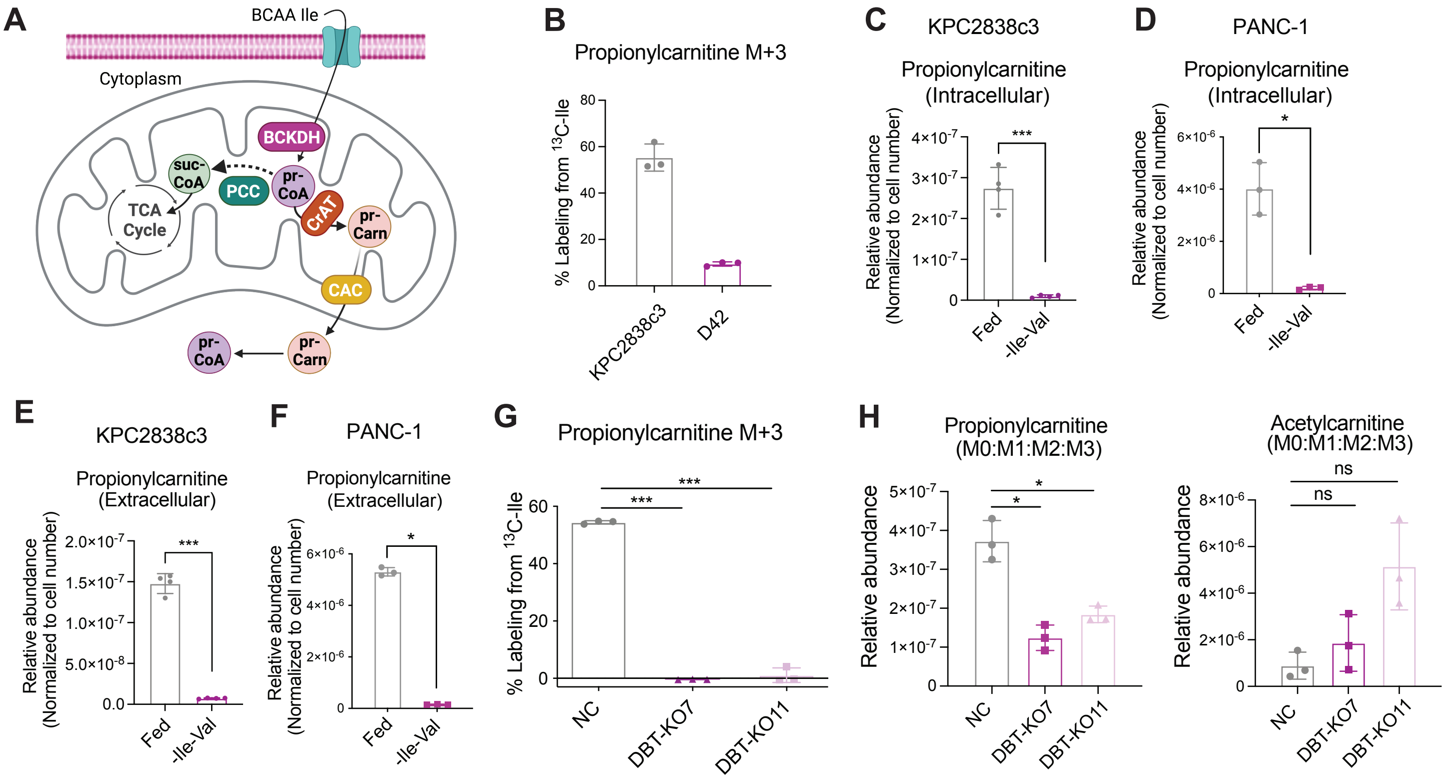

### S4

**Supplementary Figure S4**

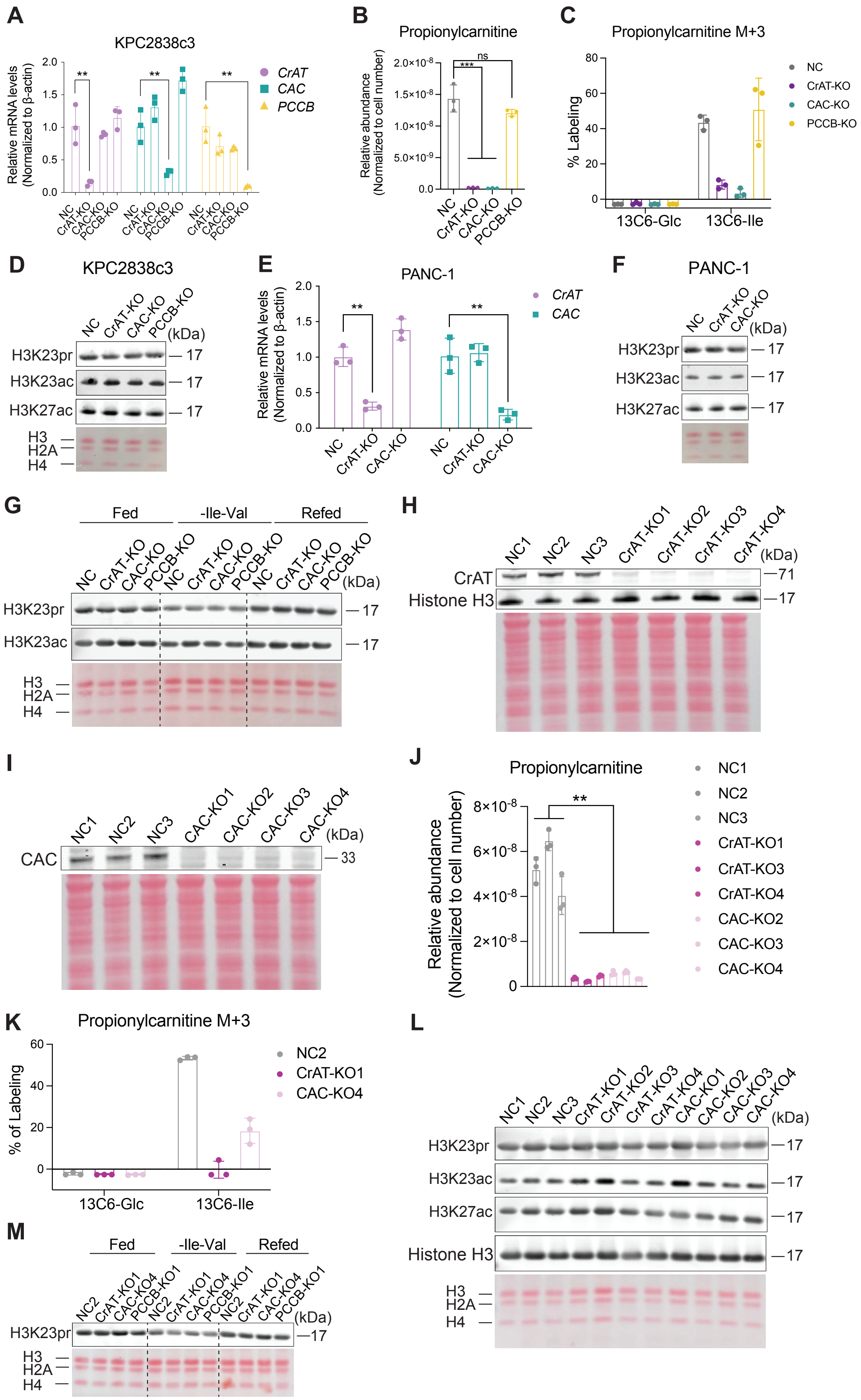

### S5

**Supplementary Figure S5**

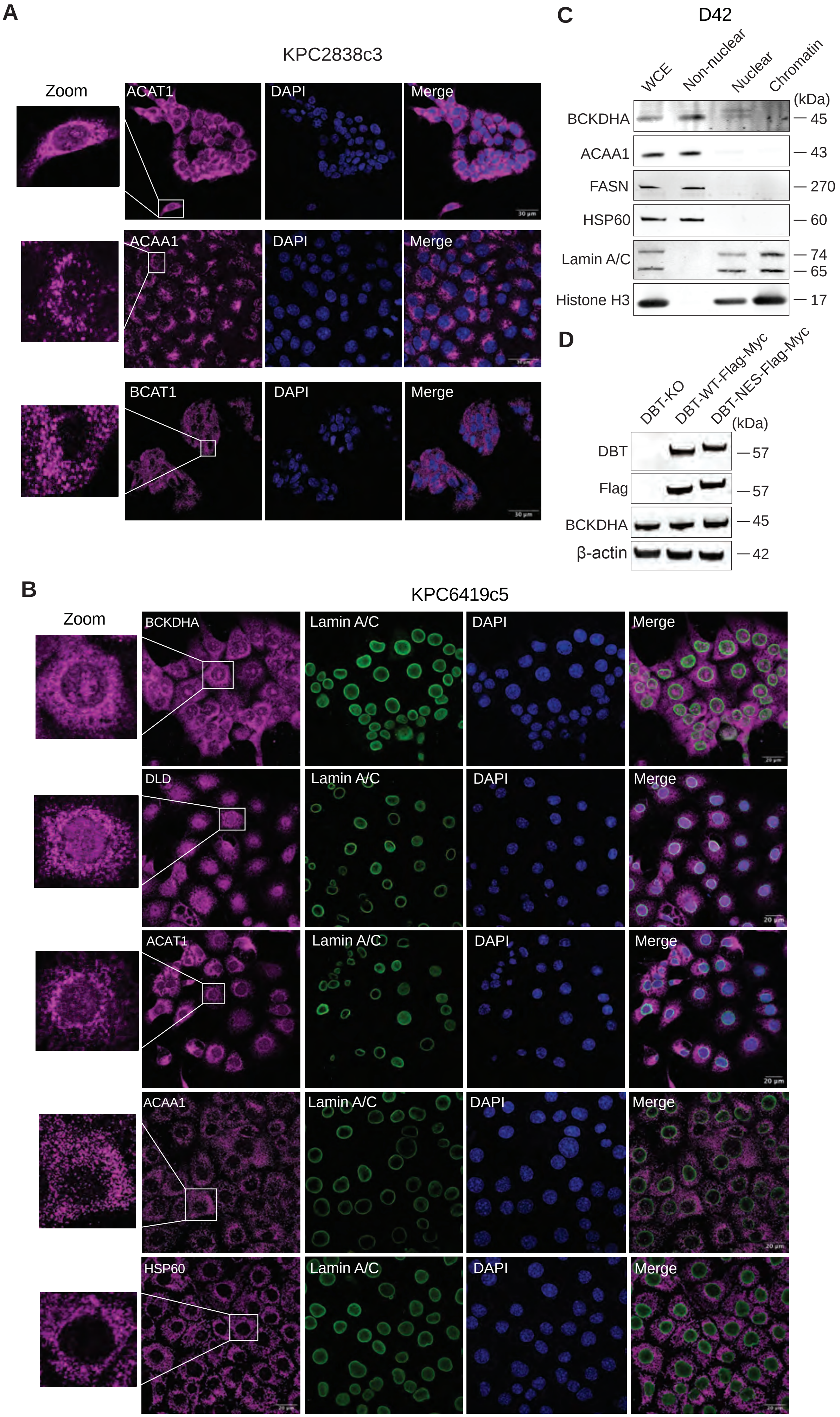

### S6

**Supplementary Figure S6**

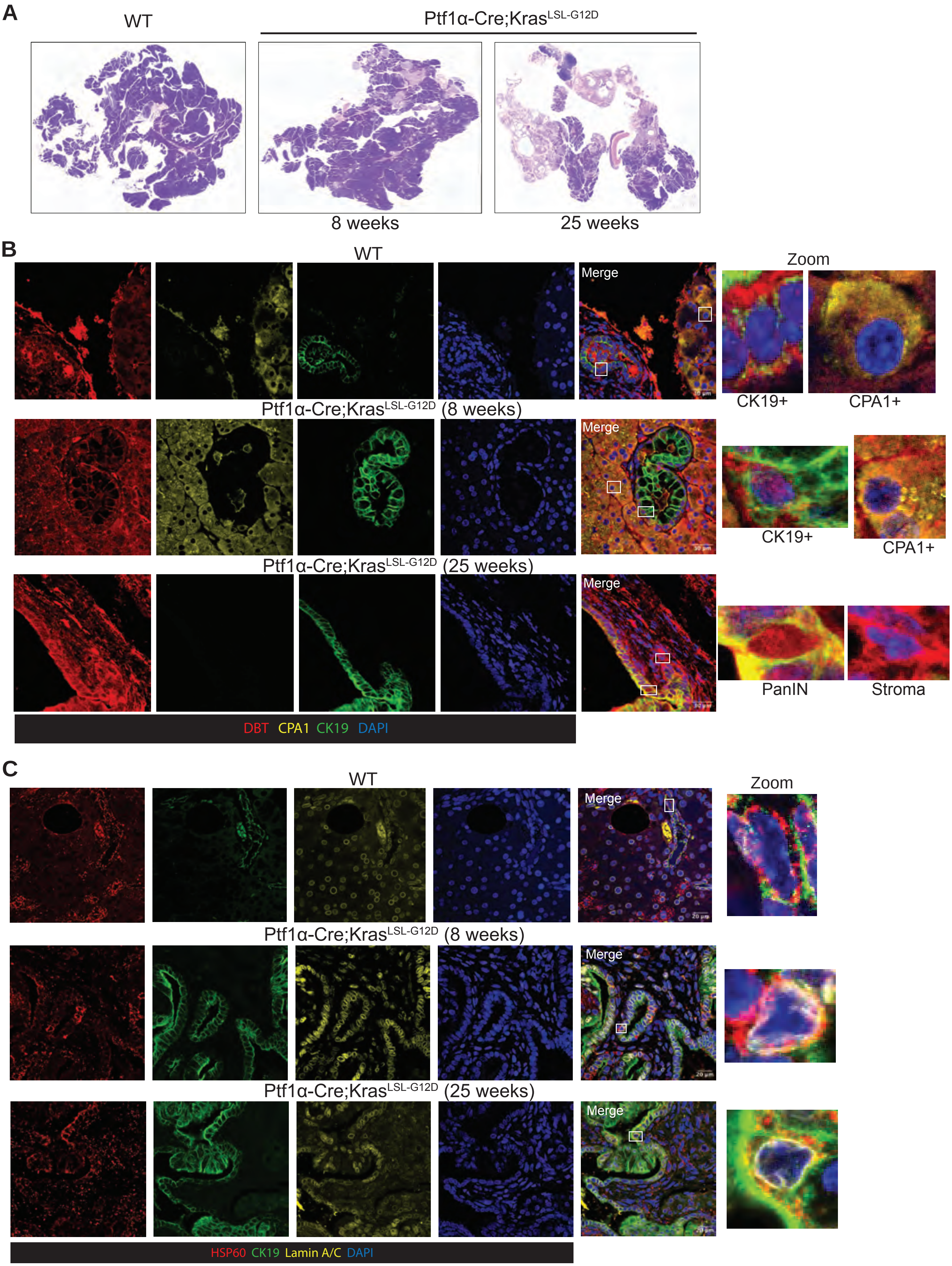

### S7

**Supplementary Figure S7**

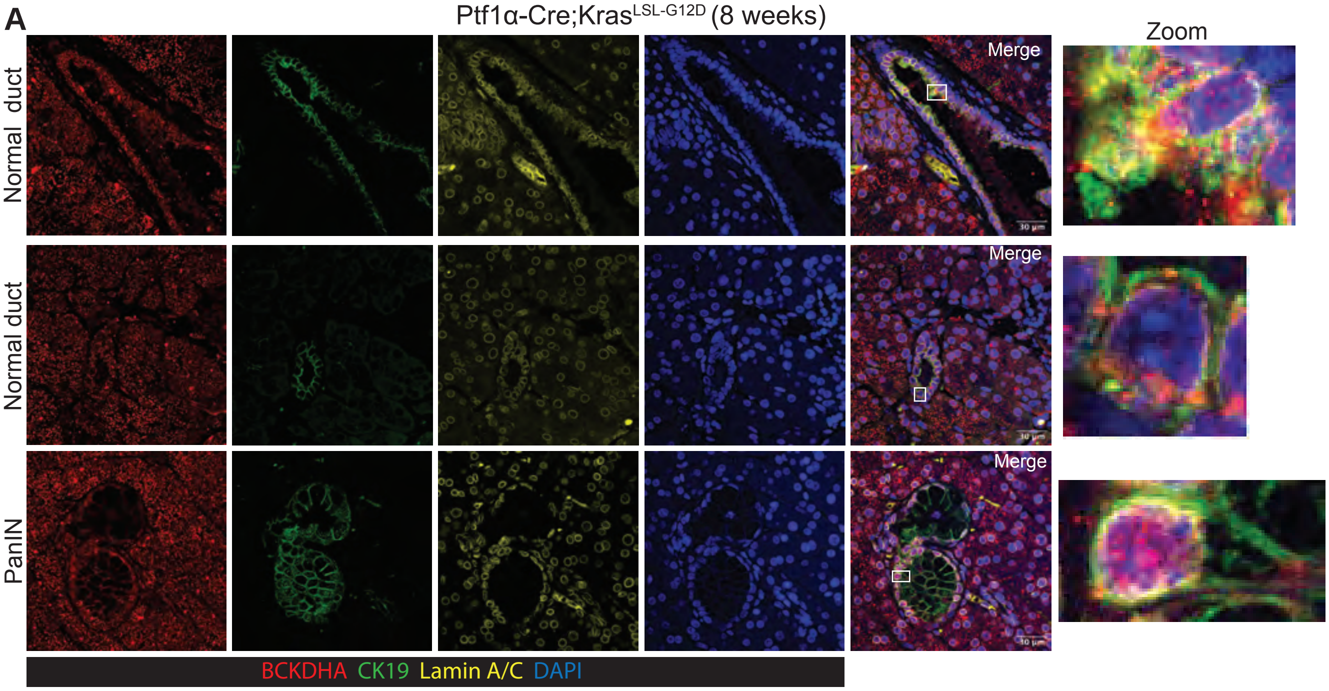


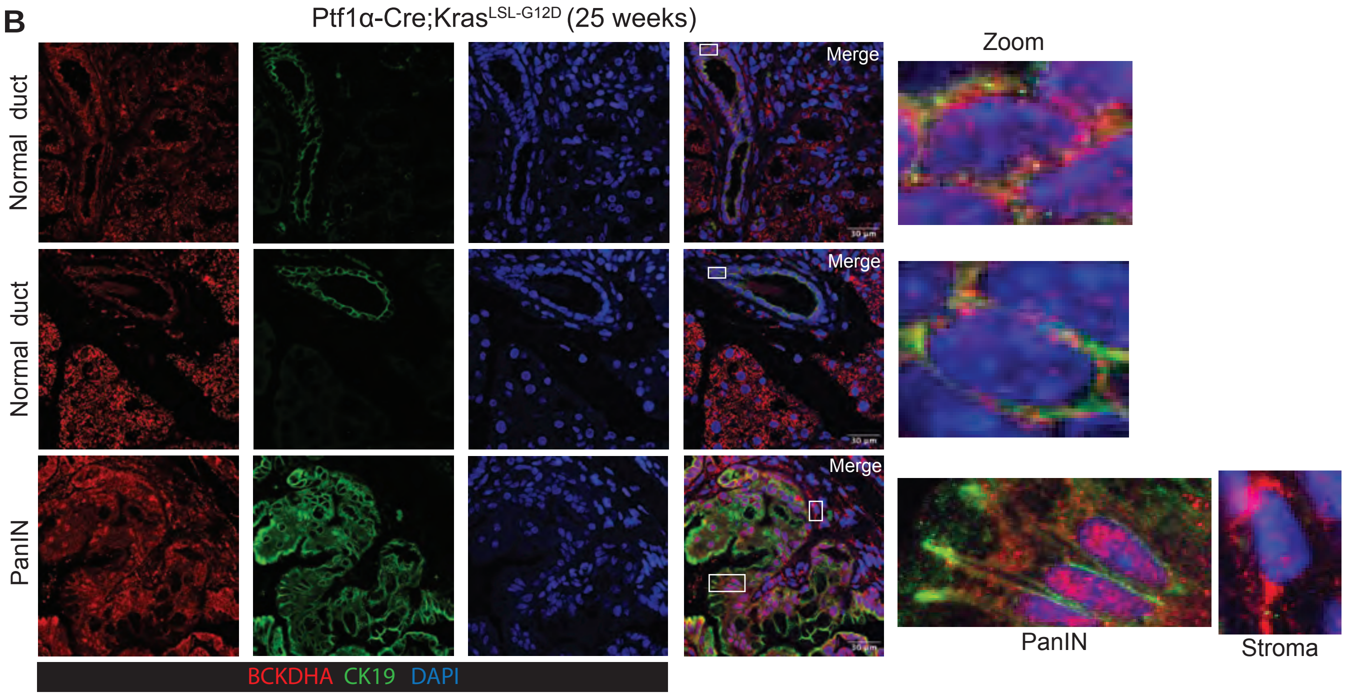

### S8

**Supplementary Figure S8**

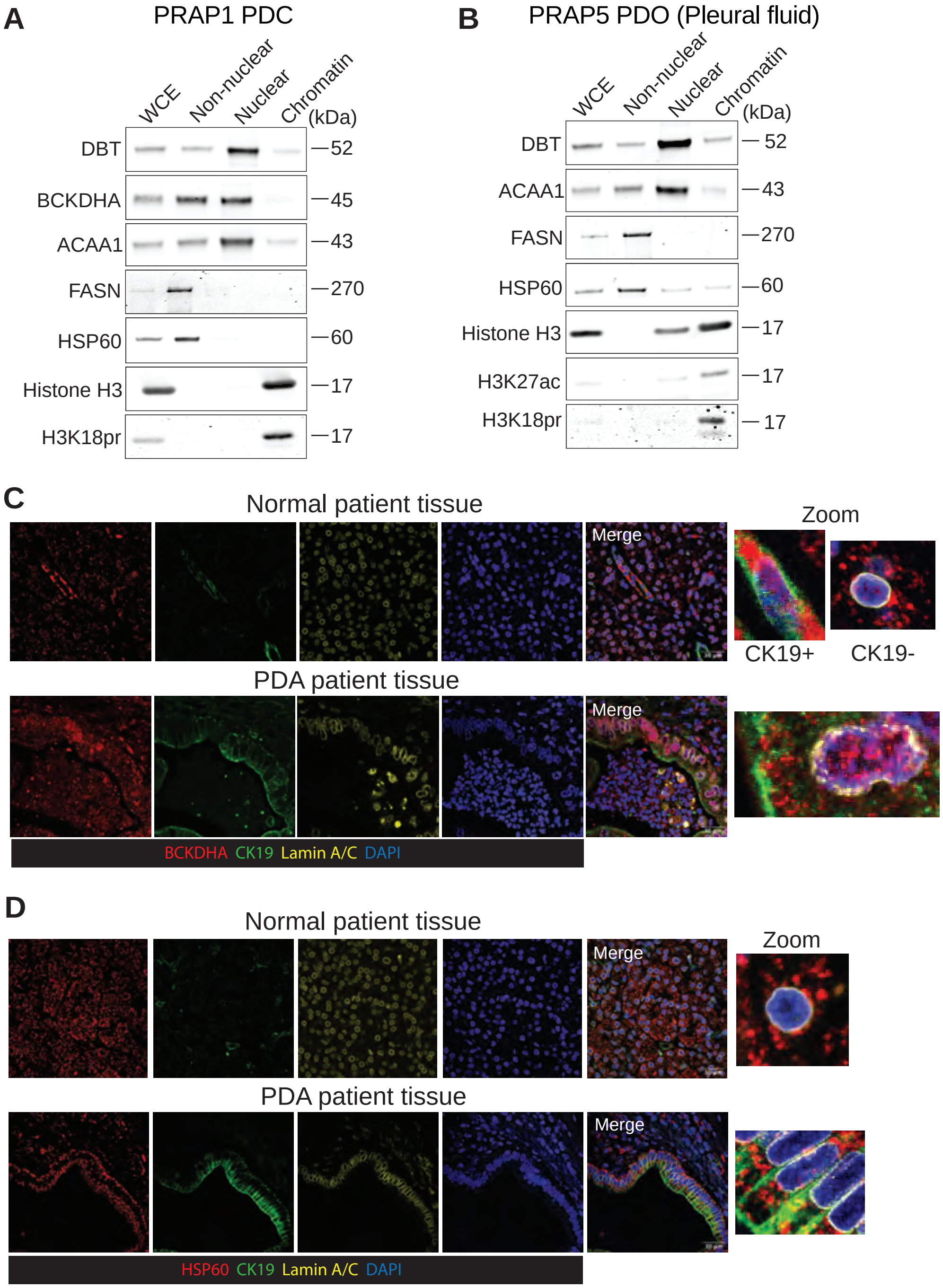

### S9

##### Supplementary Figure S9

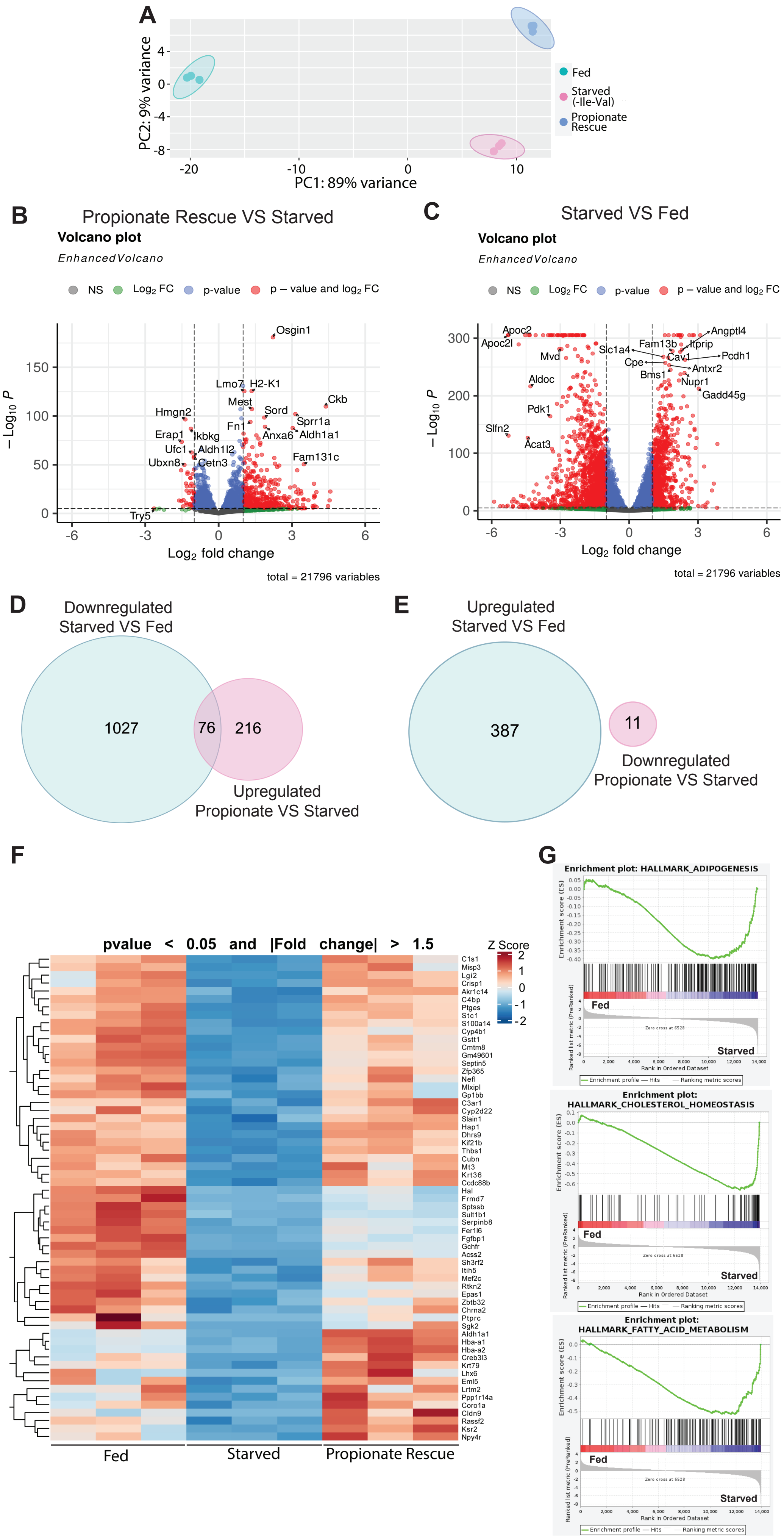

### S10

##### Supplementary Figure S10

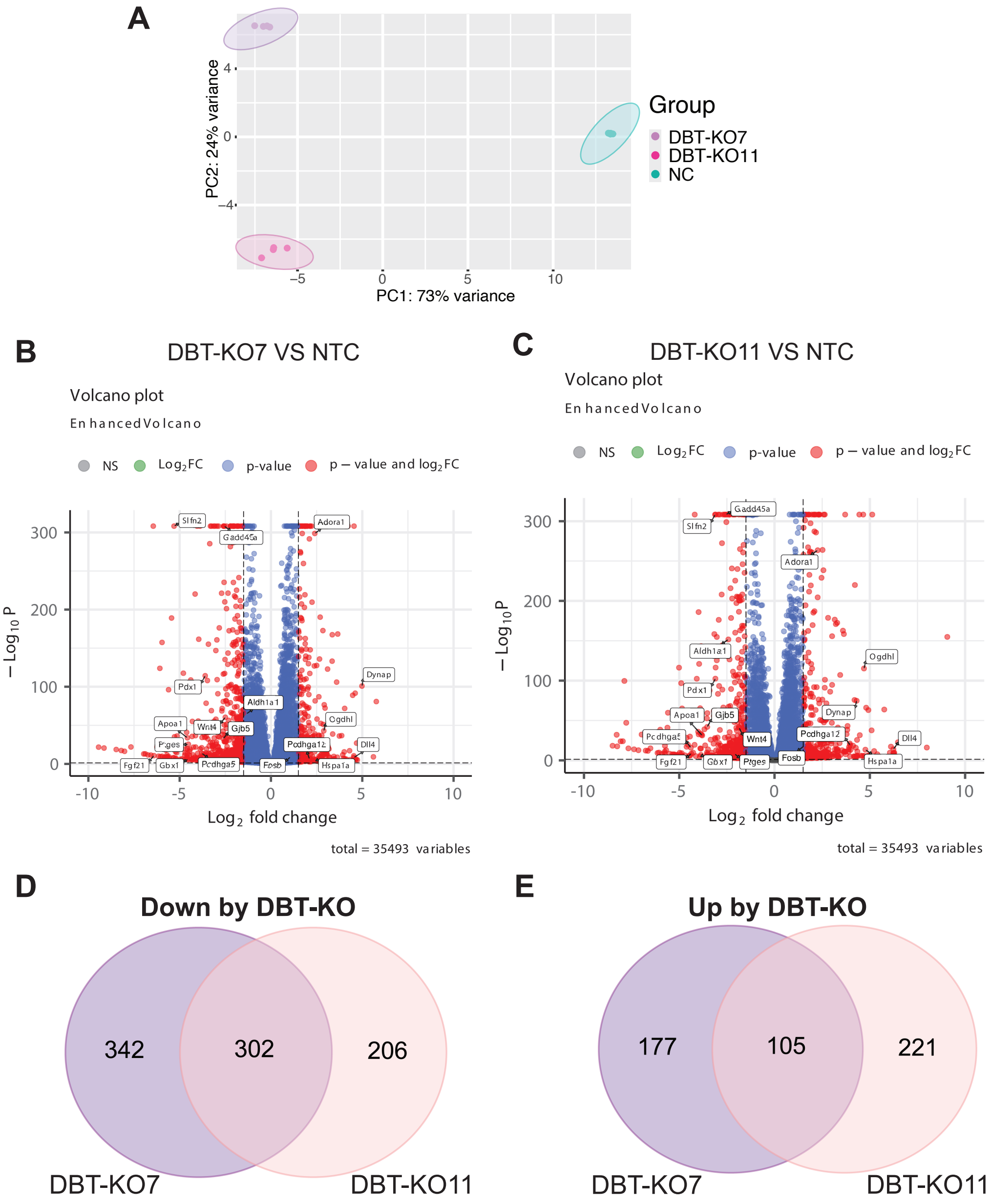

### S11

**Supplementary Figure S11**

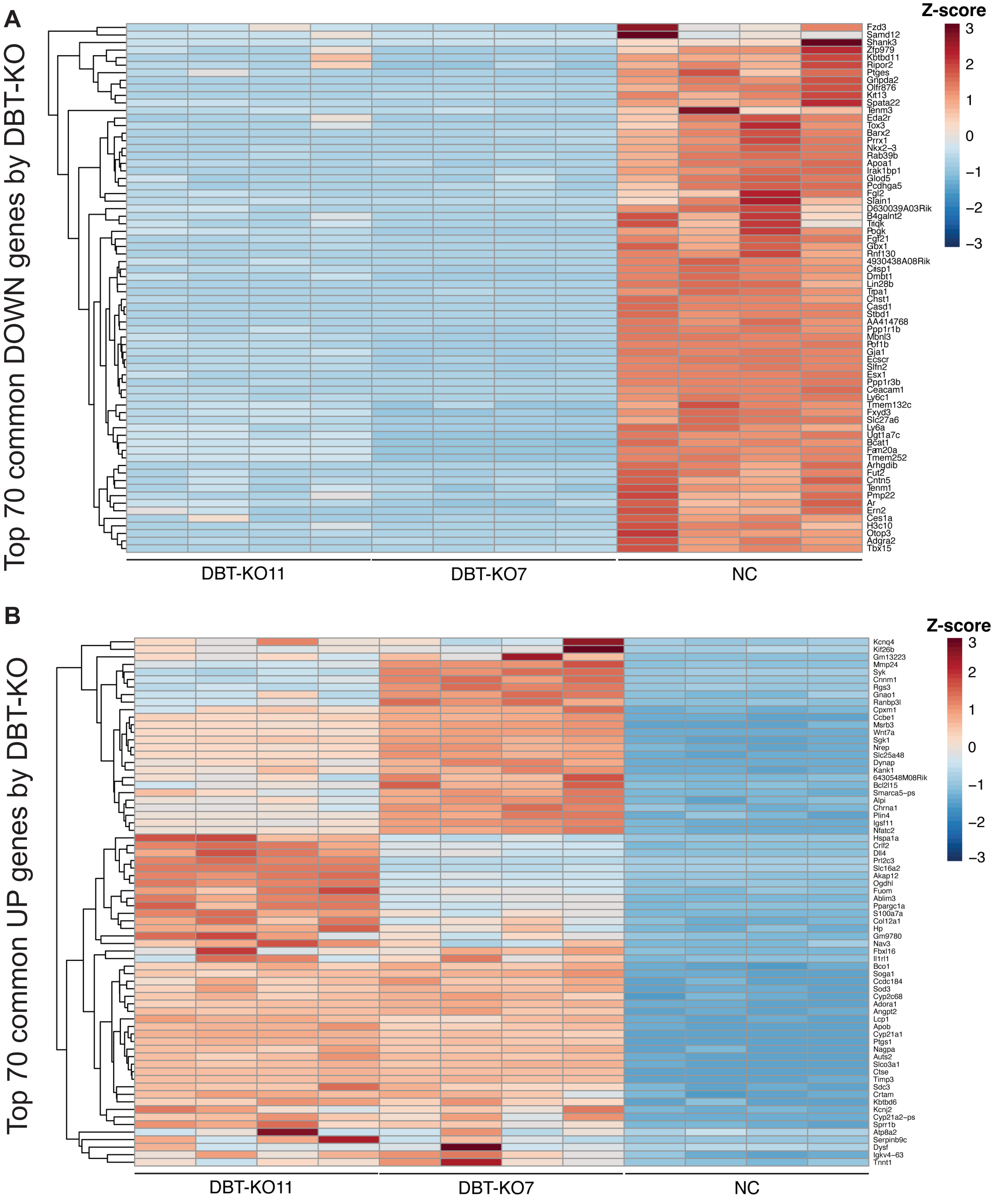

### S12

**Supplementary Figure S12**

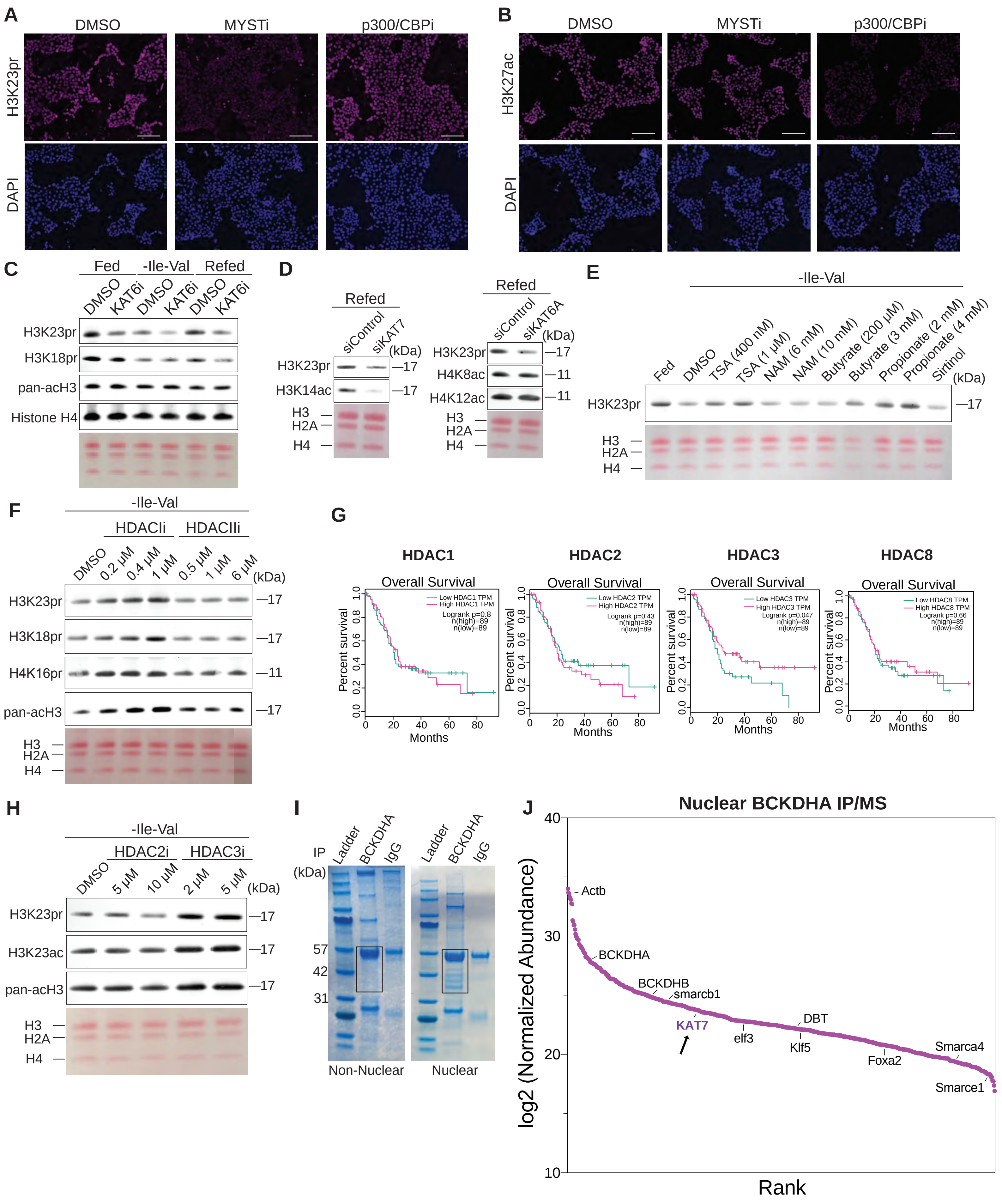
