## Supplementary figure legends for "A nuclear branched-chain amino acid catabolism pathway controls histone propionylation in pancreatic cancer"

**Supplementary Fig. S1. Histone propionylation is sensitive to BCAA catabolism in PDA.** (**A**) Proliferation of KPC2838c3 cells cultured under Fed, -Ile-Val, and -Ile-Val + Propionate conditions for 24 h. (**B**) Western blots of acid extracted histones after 24 h culture in +/- Ile and Val, +/- 2 mM propionate in KPC6419c5 cells. (**C**) Relative abundance of pr-CoA and acetyl-CoA in Fed or -Ile-Val, +/- 2 mM propionate in PANC-1 cells. (**D**) Western blots of acid extracted histones after 24 h culture in +/- Ile and Val, +/- 2 mM propionate in 786-O cells. (**E**) Immunostaining of DBT protein in NC and DBT-KO11 KPC2838c3 cells. (**F**) Acetyl-CoA abundance measured by LC-MS in NC and DBT-KO11 KPC2838c3 cells. (**G**) [U-$^{13}$C]-BCAAs (Ile, Val, Leu) were traced into acetyl-CoA M+2 in DBT-KO (clones 7 and 11) or NC control KPC2838c3 cells. (**H**) Absolute abundance of propionyl-CoA (left panel) and acetyl-CoA (right panel) in NC and BCAT2-KO KPC2838c3 whole cells. (**I**) Western blots in whole-cell extracts or acid extracted histones from NC, BCAT1-KO, and BCAT2-KO KPC2838c3 cells. (**J**) Representative images of colony formation assay using the indicated KPC2838c3 cell lines that were treated or not with 2 mM propionate.

**Supplementary Fig. S2. Impaired tumor growth correlates with elevated CD3+ T-cell infiltration.** (**A**) Representative immunofluorescence staining with Ki67 (red), Lamin A/C (yellow), and CK19 (green) antibodies in NC, DBT-KO7 and DBT-KO11 allograft tumors (n=3). Nuclei were stained with DAPI. Scale bars, 30 μm. (**B**) Representative immunofluorescence staining of CD3 protein in NC, DBT-KO7 and DBT-KO11 allograft tumors (n=3). Nuclei were stained with DAPI. Scale bars, 30 μm.

**Supplementary Fig. S3. Propionylcarnitine is sensitive to the BCAA availability. (A)** Schematic of mitochondrial pr-CoA utilization and export through the carnitine shuttle pathway. (**B**) Percent labeling of propionylcarnitine M+3 from 6 h [U-$^{13}$C]-isoleucine tracing in KPC2838c3 and D42 cells. (**C** to **D**) Intracellular levels of propionylcarnitine under fed and -Ile-Val conditions in (**C**) KPC2838c3 and (**D**) PANC-1 cells. (**E** to **F**) Extracellular abundance of propionylcarnitine from media under fed and -Ile-Val conditions in (**E**) KPC2838c3 and (**F**) PANC-1 cells. (**G**) Percent labeling of propionylcarnitine M+3 from 6 h [U-$^{13}$C]-isoleucine tracing in NC, DBT-KO7 and DBT-KO11 KPC2838c3 cell lines. (**H**) Relative abundance (sum of isotopologues) of propionylcarnitine and acetylcarnitine in NC, DBT-KO7 and DBT-KO11 KPC2838c3 cell lines. Data are presented from independent replicates. Statistical significance was determined by Student's *t*-test; error is reported as SD (*$p<0.05$, ***$p<0.001$, ns=no significance).

**Supplementary Fig. S4. The carnitine shuttle pathway is not required for BCAA sensitive histone propionylation. (A)** qPCR mRNA expression of *CrAT*, *CAC*, and *PCCB* in NC, CrAT-KO, CAC-KO and PCCB-KO KPC2838c3 cell lines. (**B**) Relative propionylcarnitine levels measured by LC-MS in the indicated cell lines. (**C**) Fraction of labeling of propionylcarnitine M+3 from [U-$^{13}$C]-isoleucine and [U-$^{13}$C]-glucose in the indicated KPC2838c3 cell lines. (**D**) Western blots of histones extracted from the NC, CrAT-KO, CAC-KO and PCCB-KO KPC2838c3 cells. (**E**) qPCR mRNA expression of *CrAT* and *CAC* in NC, CrAT-KO, and CAC-KO PANC-1 cell lines. (**F**) Western blots of histones extracted from the NC, CrAT-KO, and CAC-KO PANC-1 cells. (**G**) Western blots of histones extracted from the NC, CrAT-KO, CAC-KO and PCCB-KO KPC2838c3 cells cultured in fed, -Ile-Val and Refed conditions. (**H** to **I**) Western blot of (**H**) CrAT protein and (**I**) CAC protein in whole-cell lysates from NC (1/2/3) and KO (1/2/3/4) KPC2838c3 clonal cell lines. (**J**) Relative propionylcarnitine abundance in NC, CrAT-KO, and CAC-KO clonal cell lines. (**K**) Fraction of labeling of propionylcarnitine M+3 from [U-$^{13}$C]-isoleucine and [U-$^{13}$C]-glucose in the NC2, CrAT-KO1 and CAC-KO4 clonal cell lines. (**L**) Western blots of acid extracted histones in the indicated KPC2838c3 clonal cell lines. (**M**) Western blot of H3K23pr under Fed, -Ile-Val and Refed conditions in the indicated cell lines. Data are presented as mean ± s.d. from three independent experiments. Statistical analysis was performed using two-tailed unpaired Student's *t*-test; *p<0.05, **p<0.01, ***p<0.001, ns=no significance.

**Supplementary Fig. S5. BCAA metabolic enzymes are present in the nuclei of PDA cells. (A)** Immunostaining of ACAT1, ACAA1 and BCAT1 (pink) in KPC2838c3 cells. Nuclei were stained with DAPI. Scale bars, 30 µm. (**B**) Immunostaining of BCKDHA, DLD, ACAT1, ACAA1, HSP60 (pink) and Lamin A/C (yellow) in KPC6419c5 cells. Nuclei were stained with DAPI. Scale bars, 30 µm. (**C**) Fractionation and western blot analysis of the indicated proteins in the HCC D42 cells. (**D**) Western blot of DBT, Flag, BCKDHA and β-actin proteins in whole cell protein extracts from DBT-KO11 cells and DBT-KO11 reconstituted with DBT-WT-Flag-Myc addback and DBT-NES-Flag-Myc addback.

**Supplementary Fig. S6. Nuclear BCKDH is present in PANIN lesions.** (**A**) H&E staining of the pancreas tissue from WT C57Bl/6 mice and Ptf1α-Cre;Kras$^{LSL\text{-}G12D}$ mice at 8 weeks or 25 weeks of age. (**B**) Immunofluorescence staining of DBT (red), CPA1 acinar cell marker (yellow), and CK19 ductal cell marker (green) in the pancreas of WT, 8 weeks old Ptf1α-Cre;Kras$^{LSL\text{-}G12D}$, and 25 weeks Ptf1α-Cre;Kras$^{LSL\text{-}G12D}$ old C57Bl/6 mice. Stromal cells are negative of CK19 and CPA1. Scale bars, 30 µm. (**C**) Immunofluorescence staining of HSP60 (red), ductal marker CK19 (green) and Lamin A/C (yellow) in the pancreas tissue from WT, 8 weeks old Ptf1α-Cre;Kras$^{LSL-G12D}$, and 25 weeks Ptf1α-Cre;Kras$^{LSL-G12D}$ old C57Bl/6 mice. Scale bars, 30 μm.

**Supplementary Fig. S7. BCKDH localizes in the nuclei of PanINs but not in those of normal ducts in Kras mutant pancreas. (A)** Representative immunofluorescence images of BCKDHA (red), CK19 ductal cell marker (green), and Lamin A/C (yellow) proteins in PanIN lesions and adjacent normal ducts in the pancreas tissue of 8-weeks-old Ptf1α-Cre;Kras$^{LSL-G12D}$ mice. Scale bars, 30 μm. (**B**) Representative immunofluorescence images of BCKDHA (red) and CK19 ductal cell marker (green) in PanIN lesions and adjacent normal ducts in the pancreas tissue of 25-weeks-old Ptf1α-Cre;Kras$^{LSL-G12D}$ mice. Scale bars, 30 μm. Nuclei were stained with DAPI.

**Supplementary Fig. S8. BCAA metabolic enzymes localize in the nuclei of PDA patient cells.** (**A** to **B**) Subcellular fractionation and western blot analysis of (**A**) PRAP1 patient-derived cells (PDC) and (**B**) PRAP5 patient-derived organoids (PDO) using antibodies against the indicated proteins. Whole cell extract (WCE) was used as a control. (**C** to **D**) Representative immunostaining images of (**C**) BCKDHA (red) and (**D**) HSP60 (red) in normal and PDAC clinical samples. At least 6 areas were imaged in each individual tissue. The CK19 ductal cell marker is shown in green and Lamin A/C control is indicated in yellow color. Nuclei were stained with DAPI (blue). Scale bars, 30 μm.

**Supplementary Fig. S9. BCAA restriction alters the expression of specific genes.** (**A**) PCA plot of the transcriptome of fed (complete media with dialyzed FBS), starved (-Ile-Val), and propionate rescue (-Ile-Val+Propionate) KPC2838c3 cells from three independent biological replicates. (**B** and **C**) Volcano plots showing the differentially expressed genes in (**B**) Propionate Rescue vs Starved cells and (**C**) Starved vs Fed cells. (**D** and **E**) Venn diagrams of differentially expressed genes that are (**D**) downregulated in starvation and upregulated by propionate treatment and (**E**) upregulated in starvation and downregulated by propionate. (**F**) Summary of repressed genes by BCAA starvation which are restored by propionate rescue. (**G**) Top GSEA signatures of downregulated genes by BCAA starvation compared to fed condition.

**Supplementary Fig. S10. BCAA catabolism controls the transcription of specific genes.** (**A**) PCA plot of the transcriptome from NC (n=4), DBT-KO7 (n=4) and DBT-KO11 (n=4) KPC2838c3 cells. (**B** and **C**) Volcano plots showing the differentially expressed genes in (**B**) DBT-KO7 vs Starved cells and (**C**) DBT-KO11 vs Starved cells. (**D** and **E**) Venn diagrams showing the overlap between (**D**) downregulated and (**E**) upregulated genes in DBT-KO7 and DBT-KO11 cell lines.

**Supplementary Fig. S11. DBT loss alters the expression of selective genes.** (**A** and **B**) Heatmaps of the top 70 significantly (**A**) downregulated and (**B**) upregulated genes that are common between DBT-KO7 and DBT-KO11 compared to the NC control.

**Supplementary Fig. S12. BCAA-derived Kpr is epigenetically regulated.** (**A** to **B**) Immunostaining of (**A**) H3K23pr and (**B**) H3K27ac in KPC2838c3 cells that were starved of Ile/Val for 24 h followed by refeeding for 6 h in the presence of the indicated inhibitors. Nuclei were stained with DAPI. (**C**) Western blot of extracted histones from Fed, -Ile-Val (24h), and Refed (16h) KPC2838c3 cells that were treated with the KAT6 inhibitor WM-1119 (2 μM) or DMSO control using antibodies against the indicated marks. (**D**) Western blot of histones isolated from BCAA-refed KPC2838c3 cells treated with siC, siKAT7, or siKAT6A siRNAs for 48 h. (**E** to **F**) Western blot of histones isolated from Ile/Val-starved KPC2838c3 cells that were treated with the indicated inhibitors in **(E)** and the HDACI (Entinostat) and HDACII (MC1568) inhibitors in **(F)** or DMSO control for 24 h. (**G**) Kaplan-Meier curve showing the percent overall survival of PDA patients expressing low or high mRNA levels of HDAC1/2/3/8 using the Gepia data portal. (**H**) Western blot of histones extracted from Ile/Val-starved KPC2838c3 cells that were treated with DMSO control or inhibitors targeting HDAC2 (CAY-10683) or HDAC3 (RGFP966) using antibodies against the indicated marks. (**I**) Immunoprecipitation of BCKDHA or IgG control in the non-nuclear and nuclear fractions of KPC2838c3 cells that were ran on SDS-page gels and stained with Coomassie. The black square demonstrates the area of unique proteins precipitated with BCKDHA in the nuclear fraction that were absent from the non-nuclear fraction. (**J**) Mass-spectrometry analysis of the nuclear proteins denoted with the black square in panel (**I**). Abundance values were normalized to the total protein abundances of each sample.
