## Supplementary Table 1 for "A nuclear branched-chain amino acid catabolism pathway controls histone propionylation in pancreatic cancer"

**Table S1: Differentially regulated genes**

List of genes downregulated by starvation and rescued by propionate

Rassf2
Krt79
Kif21b
Coro1a
Akr1c14
Hba-a1
Hal
C1s1
Gm49601
Lrtm2
Syt16
Serpinb8
Ksr2
Fer1l6
Acss2
Gchfr
C3ar1
Thbs1
2210011K15Rik
Fgfbp1
Itih5
Chrna2
Eml5
Lrrc15
Misp3
Ptges
Slain1
Nefl
Sh3rf2
Hap1
Gm16201
Dhrs9
Ppp1r14a
Lhx6
Ptprc
Grhl2
Sult1b1
Crisp1
Prr15l
Cyp4b1
Gp1bb
S100a14

Gm2379
Gm19696
Lgi2
Gm4065
Rtkn2
Aldh1a1
Krt36
Septin5
Cyp2d22
Npy4r
Gstt1
Cmtm8
Tafa5
Frmd7
C4bp
Mlxipl
Vmn2r79
Mef2c
Mt3
Creb3l3
Zbtb32
Epas1
Ccdc88b
Cubn
Ms4a4d
Sgk2
Stc1
Cldn9
Hba-a2
Il13ra1
Atg9b
Cthrc1
Sptssb
Zfp365

List of common genes downregulated between Ile/Val starvation, DBT-KO7, and DBT-KO11
Selenbp1
Spdya
Myo7b
G0s2
Klb
Hba-a1
Slc16a3
Ankrd37

Mkrn2os
Syt16
Medag
Lgals4
Calcrl
Gca
Rundc3b
Fxyd3
Aldh3a1
Fancl
Cd38
Lsp1
Glod5
Fam20a
Muc4
Mfsd4a
Apoa1
Ptges
Slain1
Hif3a
Sh3rf2
Itgb2
Tmem252
Hap1
Kif12
D630039A03Rik
Fhad1
Egflam
Arhgdib
Crisp1
Kbtbd11
Slfn2
Gjb5
Krt39
Cth
Prr15l
S100a14
4930438A08Rik
Bche
B4galnt3
Fgfr4
Lgi2
Bnip3
Enpep

Clec2d
Papln
Upp1
Ceacam1
Ces1a
Vrk2
Gstt1
Spata22
Krt13
Fam83e
Smoc1
Akr1c19
Chac2
Albfm1
Ripor2
Tspan13
Ms4a4d
Ly6a
Fgl2
Tnfsf13
Cntn5
Cyp24a1
2200002J24Rik
Barx2
Ugt1a7c
Lrrc26
Clec2g
Nrg4

List of common genes between DBT-KO and propionate rescue under starvation
Prr15l
S100a14
Hba-a1
Lgi2
Syt16
Gstt1
Ptges
Slain1
Sh3rf2
Ms4a4d
Hap1
Crisp1
