## Supplementary Table 2 for "A nuclear branched-chain amino acid catabolism pathway controls histone propionylation in pancreatic cancer"

**Supplementary Table S2. Primers and oligos**

| qRT-PCR primers | | |
|---|---|---|
| **Gene name** | **Forward (5→3)** | **Reverse (5→3)** |
| mCrAT | TGGTCATCTACTCCAGCCCA | AACTGGCAGCGTCTCATTGT |
| hCrAT | GCTGCCAGAACCGTGGTAAA | CCTTGAGGTAATAGTCCAGGGA |
| mCAC | GCCTCCTGGGATAAGTGTGA | GTGCTCTCATGGGGAAAGAA |
| hCAC | GGGGTCACTCCCATGTTTG | TGTGGTGAATACGCCAGATAAC |
| PCCB | TACTCCCCTGCCCTAACAGA | GGGCCAGTGATAAACAGGTAAG |
| Ptges | GGATGCGCTGAAACGTGGA | CAGGAATGAGTACACGAAGCC |
| S100a14 | TGTAGAGAGGGCCATTGAGA | CAGGGGTCAGTGTTTCCTTT |
| Ripor2 | CTCAGGCCAAGCTGAAGAAA | AGGCTCCTTGGGAGTGTTGT |
| Lgi2 | ACCAAGGAGTCCATCATCTGC | AGGGAGCTGATGTCGCCA |
| Sh3rf2 | CACCTTCTGCAAACCATGTC | ACACCAGAGTCCTGCATTCC |
| Gstt1 | GTTCTGGAGCTGTACCTGGATC | AGGAACCTTATACTTGTGTGCC |
| Hif3a | GGACGCCTGCTACCTGAAG | GTAAGCCATGTCTCCCTCGG |
| CS | TCCGAGGCTACAGTATCCCT | CTCAGGCAGGGGTTCTTCTC |
| Nf1 | GCTTTGCTGGTCCTTCATCAG | CTACAGGAGCGTCAGGATTCC |
| mβ-actin | TGGTGGGAATGGGTCAGAA | TCTCCATGTCGTCCCAGTTG |
| hβ-actin | AGAGCTACGAGCTGCCTGAC | AGCACTGTGTTGGCGTACAG |
| GAPDH | GTGTGAACGGATTTGGCCGT | TTGATGGCAACAATCTCCAC |
| ChIP primers | | |
| Ptges | ggtgtccccgagttgaagtc | GCCGCTCTCCATCACCAG |
| Sh3rf2 | CACCTTCTGCAAACCATGTC | ACACCAGAGTCCTGCATTCC |
| Ripor2 | gtcccagagcctgacgtg | ccttcagcccgactcctt |
| CS | tgtagctctctcccttcggt | TGCAGTGAGTAGAGCCATgg |
| CRISPR guides | | |
| DBT | CACCGTGTCTGAAAATAGCGAAC AC | aaacGTGTTCGCTATTTTCAGACA C |
| mCrAT | CACCGTCCACAAGTGCAACTATG GG | aaacCCCATAGTTGCACTTGTGG AC |
| hCrAT | CACCGCAGGACTTCGTGGACCTG CA | aaacTGCAGGTCCACGAAGTCCT GC |
| mCAC | CACCGGCATCACAGGGCTGTATC G | aaacCGATACAGCCCTGTGATGC C |
| hCAC | CACCGAATGGCTGCCCCTATCATC G | aaacCGATGATAGGGGCAGCCAT TC |
| PCCB | CACCGCTTGTGCTGCGCGTCGAT G | aaacCATCGACGCGCAGCACAA GC |
| NC | CACCGACTCCGGGTACTAAATGT C | aaacGACATTTAGTACCCGGAGT C |
